# A dynamical attractor computes reward prediction in the basal forebrain

**DOI:** 10.64898/2026.10.01.755868

**Authors:** Hsiao-Chen Liu, Shih-Chieh Lin

**Affiliations:** Institute of Neuroscience, National Yang Ming Chiao Tung University, Taipei, Taiwan; Brain Research Center, National Yang Ming Chiao Tung University, Taipei, Taiwan

## Abstract

Phasic ensemble bursting in subcortical neuromodulatory systems such as the basal forebrain (BF) provides a powerful format for broadcasting reward-prediction signals. These bursts encode reward prediction by varying population amplitude, yet their internal structure, reflected in cell-to-cell amplitude relationships, remains conserved. How neural circuits reconcile these competing constraints remains unknown. Here we show that BF bursting is generated by a dynamical attractor: burst structure is imposed not by feedforward inputs but by recurrent circuit dynamics, while burst amplitude is determined by approximately linear integration across concurrent inputs. Optogenetic activation of BF neurons expressing natriuretic peptide receptor 3 (Npr3) outside the behavioral task reproduced the low-dimensional dynamics of natural cue-evoked bursting, preserving the temporal envelope and cell-to-cell amplitude relationships across stimulation patterns. Critically, unilateral activation of non-overlapping Npr3 populations in either hemisphere recruited the full bilateral ensemble into the same native trajectory, showing that restricted inputs can nevertheless recruit the full ensemble. Matched ipsilateral and contralateral onset latencies exceeding 25 ms implicated recurrent interactions beyond the BF. Despite the conserved structure, population burst amplitude varied with stimulation strength and summed approximately linearly across concurrent unilateral inputs. This linear summation extended to natural sensory input during behavior: adding Npr3 stimulation to auditory cues increased cue-evoked burst amplitude, improved detection, and reduced subsequent reward responses, consistent with stronger reward prediction producing smaller reward prediction errors. Thus, a low-dimensional dynamical attractor preserves a shared BF burst structure across diverse inputs while integrating concurrent inputs into a scalar burst amplitude that encodes reward prediction.

## Introduction

Phasic population bursting provides a common format for reward-prediction signaling in subcortical neuromodulatory systems. Canonical examples include midbrain dopamine neurons^1–3^, as well as a distinct population of noncholinergic basal forebrain (BF) neurons, termed BF bursting neurons, which exhibits similarly rapid phasic bursting linked to reward prediction and behavioral performance^4–6^. In both systems, diverse sensory cues can recruit similar phasic population responses, with amplitude varying along a scalar dimension that reflects the strength of reward prediction. However, these population responses are not defined by amplitude alone: they also exhibit a reproducible internal structure, reflected in stable cell-to-cell amplitude relationships, or neuron-specific amplitude scaling^3,4,7^. This means that neurons contributing more strongly to one population burst remain relatively stronger contributors across other conditions, even as overall population burst amplitude varies (Figure 1A). How diverse inputs converge onto a shared population signal with scalable amplitude while preserving this internal structure remains unclear.

**Figure 1.**
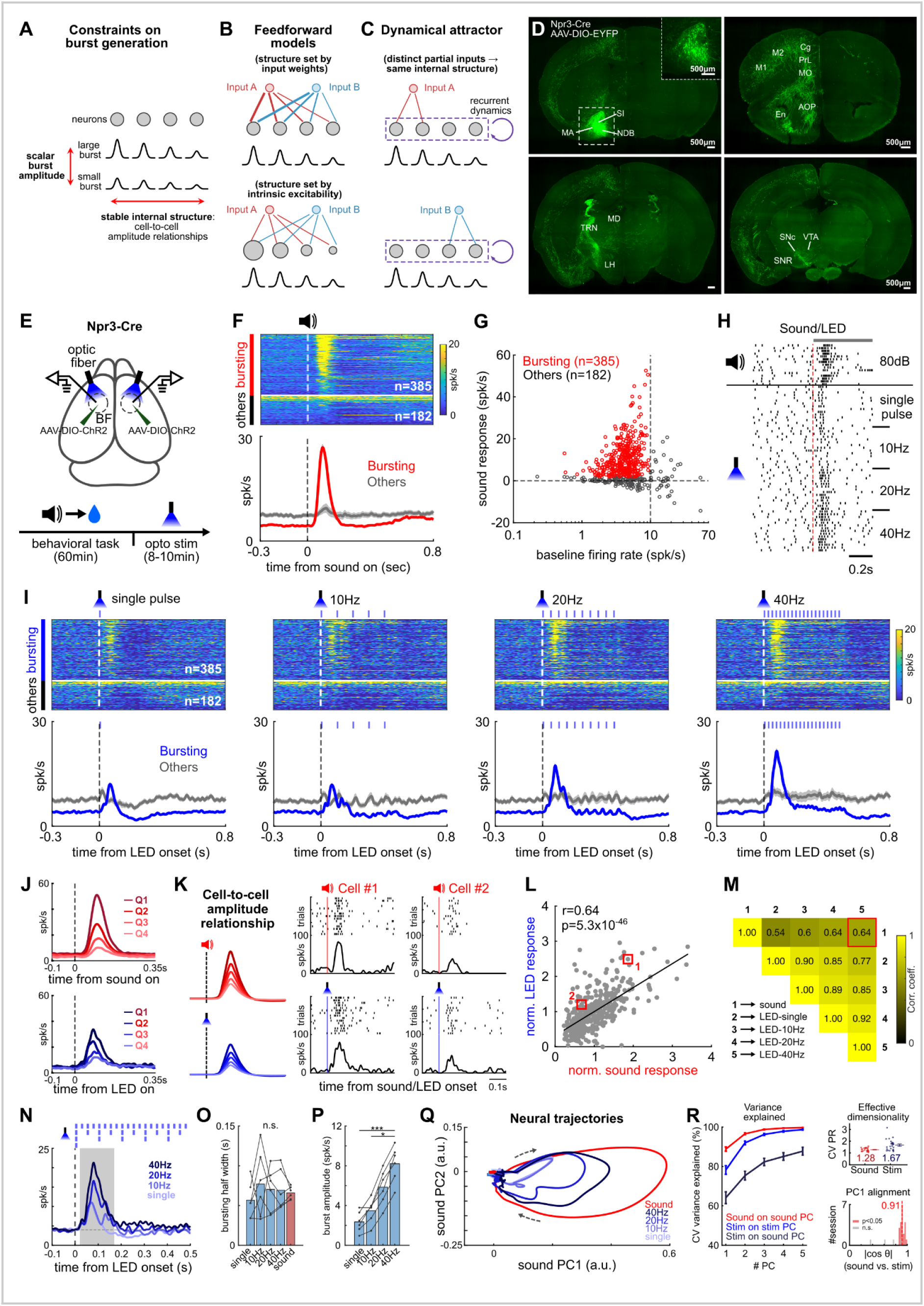
Npr3 stimulation selectively recruits the endogenous BF burst ensemble and preserves its internal structure. **A**, Two constraints on burst generation: population burst amplitude varies across events, whereas cell-to-cell amplitude relationships remain stable. **B**, Feedforward models in which stable burst structure is imposed either by matched input weights (top) or by intrinsic excitability of individual neurons (bottom). In both cases, preserving the same cell-to-cell amplitude relationships requires distinct inputs to access essentially the full ensemble. **C**, Dynamical attractor model. The preserved output pattern is generated by a recurrently connected network, allowing distinct partial inputs to recruit the full ensemble into the same structured burst. **D,** Anterograde tracing after unilateral AAV-EF1α-DIO-EYFP injection into the BF of Npr3-Cre mice, showing EYFP-expressing somata in BF regions including SI, MA, and NDB, and prominent ipsilateral axonal projections to cortex, TRN and VTA/SNc (n = 7 animals). **E,** Experimental design for simultaneous BF recording and bilateral optogenetic stimulation of Npr3 neurons. Mice first performed an auditory detection task, followed by randomly interleaved LED stimulation protocols delivered in the absence of auditory cues or rewards. **F,** Population heatmap and average PSTHs (mean±s.e.m.) showing BF responses to reward-predicting sounds for BF bursting neurons and other BF neurons. Neurons were sorted by response amplitude in the 0.05-0.2 s window after sound onset. **G,** Classification of BF bursting neurons based on strong phasic sound-evoked responses and low baseline firing rates. **H,** Example BF bursting neuron showing phasic responses to both reward-predicting sound and LED stimulation. **I,** Population heatmaps and average PSTHs (mean±s.e.m.) showing that Npr3 stimulation selectively recruited BF bursting neurons across stimulation protocols; neurons are shown in the same order as in **F**. Other BF neurons showed little response. Optogenetic stimulation patterns are indicated by blue ticks. **J,** BF bursting neurons grouped by quartiles of sound-evoked response amplitude showed similarly graded responses to 40-Hz LED stimulation. **K,** Schematic of conserved cell-to-cell response amplitude relationships during bursting, with two example neurons illustrating preserved relative amplitudes during sound-evoked and LED-evoked responses. **L,** Conserved cell-to-cell amplitude relationships between sound-evoked and LED-evoked BF bursting responses, shown by Pearson correlation of normalized response amplitudes across individual BF bursting neurons. Response amplitudes were normalized by the average response amplitude of all simultaneously recorded BF bursting neurons. Boxed points indicate the example neurons shown in **K**. **M,** Correlation matrix of normalized response amplitudes across sound and LED stimulation conditions. All pairwise correlations were significant (Pearson correlation, all p < 10^−29^). **N,** Average BF bursting responses evoked by single-pulse, 10 Hz, 20 Hz, and 40 Hz LED stimulation. Optogenetic stimulation patterns are indicated by blue ticks. Despite continued stimulation during pulse trains, BF activity remained dominated by a single early burst. **O,** Response half-widths of sound-evoked and LED-evoked BF bursting did not differ across conditions (Friedman test, p > 0.05). **P,** Mean firing rates of LED-evoked bursting responses increased in higher frequency stimulation protocols (Friedman test followed by Bonferroni-corrected pairwise Wilcoxon signed-rank tests). Each dot in **O** and **P** represents one animal, averaged across all contributing sessions (n = 6 animals). n.s., not significant; *p < 0.05, **p < 0.01, ***p < 0.001. **Q,** Population neural trajectories. Mean sound (red) and stimulation (blue) responses projected onto the first two PCs of the sound response. Arrows indicate progression over time from baseline through the burst and back to baseline. **R,** Left: Cross-validated cumulative variance explained by the first *k* PCs for sound on sound PCs (red), stimulation on stimulation PCs (blue; ceiling), and stimulation on sound PCs (black; alignment). Mean ± s.e.m. across 25 sessions. Right, top: Effective dimensionality, quantified by cross-validated participation ratio (CV PR), for sound and stimulation; dots, sessions; mean ± s.e.m., with group means shown below. Right, bottom: PC1 alignment across sessions, quantified as |cos θ| between sound and stimulation PC1 loading vectors (1, identical; 0, orthogonal). Filled bars, *p* < 0.05; open bars, n.s.; dashed red line, median.

A conventional explanation is that phasic population bursting is generated by feedforward drive. Under this account, the preserved internal structure could arise either from distinct inputs carrying matched neuron-specific weights or from stable differences in intrinsic excitability among the bursting neurons (Figure 1B). In the first case, the input specifies the relative response amplitudes directly; in the second, the same relative amplitudes emerge from properties of the output neurons themselves. While these mechanisms are not mutually exclusive, any such direct-drive account requires all effective inputs to access essentially the full bursting ensemble, regardless of whether the inputs arise from different sensory modalities, stimulus features, or reward-related events. It is not obvious that such anatomically and functionally diverse inputs would each have broad access to the same distributed bursting population. If, on the other hand, an input reaches only a subset of the circuit, how could it nevertheless recruit the full population burst while preserving its internal structure and still allow overall population burst amplitude to vary along a scalar dimension?

One possible solution is that the shared output format is generated by an internally organized dynamical attractor rather than imposed directly by feedforward input (Figure 1C). In an attractor framework, diverse inputs can act as triggers that drive the network into the same internally generated population trajectory, whose structure is determined by recurrent circuit dynamics rather than by the detailed pattern of the initiating input. This provides a direct solution to the access problem: an input that reaches only a subset of the circuit could nevertheless recruit the full structured population burst. Importantly, this framework may also accommodate a scalar amplitude dimension: population activity can be constrained to a shared low-dimensional trajectory while specific output dimensions, such as population burst amplitude, remain free to vary with input drive^8–11^. Once initiated, the trajectory should evolve according to internal circuit dynamics rather than simply track the ongoing temporal structure of the input. Whether such an internally organized dynamic can account for phasic reward-prediction bursting in a subcortical neuromodulatory system has not been tested.

The BF provides a tractable system for testing this possibility because distinct reward-predicting cues are known to converge onto the same BF bursting neurons while preserving their cell-to-cell amplitude relationships^4,7^. A key test of the dynamical attractor hypothesis is whether an input that accesses only a subset of the circuit can nevertheless recruit the full structured burst. Guided by recent single-cell and spatial transcriptomic atlases that resolve the molecular heterogeneity of the BF and adjacent pallidal territories^12–15^, we identify BF Npr3 (natriuretic peptide receptor 3)-expressing GABAergic neurons as an experimental entry point that can trigger BF bursting outside the behavioral task. Using this entry point, we asked whether unilateral Npr3 activation could recruit the full structured burst bilaterally, a key prediction of the dynamical attractor model. We then asked how population burst amplitude is determined when multiple inputs are present simultaneously, and whether the same integration principle extends to natural reward-predicting input during behavior. Finally, we asked whether Npr3-induced BF bursting predicts behavioral performance and subsequent reward responses in the same manner as natural BF bursting. Together, these experiments test whether a restricted entry population can recruit a full internally organized BF burst whose scalar amplitude integrates concurrent inputs and encodes reward prediction to modulate behavioral performance.

## Results

### Npr3 identifies a distinct GABAergic population within the basal forebrain

To test whether BF ensemble bursting reflects an internally organized burst mode, we needed an experimentally accessible entry point for recruiting BF bursting neurons, which have been studied primarily in the substantia innominata (SI) region^4–7^. Such an entry point required a molecularly defined population anatomically embedded within this BF territory. Although the BF is best known for its cholinergic neurons, it also contains prominent noncholinergic populations, including GABAergic neurons^16–19^. Within the SI, these GABAergic neurons are themselves highly heterogeneous, reflecting the anatomical complexity of a region where multiple macrosystems intersect and intermingle, particularly around the posterior part of the ventral pallidum (VP)^15,20,21^. Recent single-cell and spatial transcriptomic atlases provide a systematic framework for resolving this heterogeneity and for evaluating candidate marker genes in relation to their anatomical distributions across cell types^12–15^. Using this approach, we selected Npr3 as a candidate marker for a distinct GABAergic population because its spatial expression pattern closely mirrored the anatomical organization of the broader BF macrosystem across its anteroposterior extent, from the medial septum and diagonal band through the SI and magnocellular preoptic area to the external globus pallidus (GPe)^16,22^ (Figure S1). Within subcortical regions, Npr3 expression was highly selective for these BF-associated territories and sparse in surrounding structures, providing a clear anatomical contrast with neighboring regions (Figure S1).

We next characterized the projection targets of Npr3 neurons in the BF using cell-type-specific anterograde tracing in Npr3-Cre mice, with viral labeling targeted to the SI and adjacent BF territories (Figure 1D). Npr3 axons densely innervated deep layers of frontal cortex, the thalamic reticular nucleus, and midbrain regions including the ventral tegmental area (VTA) and substantia nigra pars compacta (SNc) (Figure 1D), recapitulating a subset of projection targets previously described for BF GABAergic neurons^19^. Npr3+ neurons in the neighboring GPe similarly project to the cortex, thalamic reticular nucleus and SNc^23,24^. By contrast, BF Npr3 neurons did not project to canonical VP output targets, such as thalamic relay nuclei or substantia nigra pars reticulata, and lacked projections to the contralateral BF (Figure 1D). Thus, Npr3 defines a distinct GABAergic projection population associated with the broader BF macrosystem.

### Npr3 stimulation selectively recruits the endogenous BF bursting ensemble with preserved cell-to-cell amplitude relationships

Having identified Npr3 neurons as a genetically defined GABAergic population within the broader BF macrosystem, we next asked whether this population could provide an experimental entry point for activating BF ensemble bursting outside a behavioral context while reproducing the structure of natural cue-evoked bursting. To test this, we first used an auditory task in which a clearly audible sound predicted reward, allowing reliable identification of BF bursting neurons according to established criteria (Figure 1E-H; Figure S2). We then activated Npr3 neurons optogenetically outside the task and in the absence of reward (Figure 1E), allowing us to test whether Npr3 input could recruit the endogenous BF burst mode independently of the behavioral events that normally evoke it.

BF Npr3 neurons in Npr3-Cre mice were selectively labeled with ChR2 using a Cre-dependent viral vector and optogenetically activated with 465-nm light pulses (10 ms, 4-6 mW), delivered either singly or as 500-ms trains at frequencies up to 40 Hz. Directly activated, phototagged Npr3 neurons were rare in our recordings (Figure S3). Nevertheless, Npr3 activation produced robust BF responses, indicating substantial indirect recruitment of the broader BF population. BF bursting neurons were identified independently based on their characteristic phasic responses to the reward-predicting sound according to established criteria^4–7,25,26^ (Figure 1F,G). Across stimulation protocols, Npr3 activation preferentially recruited BF bursting neurons with a phasic response (Figure 1H,I), while other BF neurons showed weaker and more heterogeneous modulation. BF bursting neurons grouped into quartiles by sound-evoked response amplitude showed correspondingly graded responses across Npr3 stimulation protocols (Figure 1J). At the single-neuron level, BF bursting neurons with relatively stronger sound-evoked responses also showed relatively stronger Npr3-evoked responses, preserving cell-to-cell amplitude relationships across conditions (Figure 1K-M; Pearson correlation, r = 0.54-0.92). Notably, this structured recruitment was evident even after a single stimulation pulse, showing that transient Npr3 activation can potently trigger the endogenous structured BF burst mode.

### BF burst temporal structure is conserved across Npr3 input patterns while population burst amplitude scales with input strength

We next asked whether the temporal structure of the evoked burst was dictated by the stimulation pattern or constrained by the dynamics of the circuit generating BF bursting. Despite large differences in the temporal pattern of stimulation, BF responses converged onto a similar temporal burst envelope (Figure 1N). On the one hand, a single 10-ms pulse evoked a prominent BF burst that substantially outlasted the stimulation itself, indicating that Npr3 activation did not merely produce a transient input-locked response. On the other hand, 500-ms pulse trains extending well beyond the BF burst window did not produce proportionally prolonged responses. Indeed, burst durations did not differ significantly across natural sound-evoked bursting and the single-pulse, 10-Hz, 20-Hz, and 40-Hz Npr3 stimulation conditions (Friedman test, p > 0.05; Figure 1O). Thus, the temporal envelope of the BF burst was constrained by circuit dynamics rather than by the stimulation pattern. Once the burst trajectory was initiated, later pulses exerted progressively reduced effects, revealing nonlinear, state-dependent sensitivity to additional input. This nonlinear state dependence and conserved temporal architecture are hallmark features of dynamical attractor behavior.

Despite this constrained temporal evolution, population burst amplitude remained graded with input strength. Population burst amplitude increased systematically across single-pulse, 10-Hz, 20-Hz, and 40-Hz stimulation conditions (Friedman test followed by Bonferroni-corrected pairwise Wilcoxon signed-rank tests, *p* < 0.05; Figure 1P), indicating that burst amplitude provides a scalable output dimension that preserves the ordering of input strength.

### Npr3 stimulation recruits the low-dimensional dynamics of natural BF bursting

We next asked whether Npr3-evoked bursting shared the low-dimensional population dynamics of natural BF bursting. Principal component analysis (PCA) revealed that natural sound-evoked and Npr3-evoked responses followed closely aligned population trajectories, differing primarily in the magnitude of their excursion rather than in trajectory direction or shape (Figure 1Q). Cross-validated PCA^27^ confirmed that both responses were strongly low-dimensional: PC1 alone accounted for 88.8% and 78.2% of the reproducible population variance for sound- and Npr3-evoked activity, respectively. Effective dimensionality, quantified by the participation ratio^28^, was 1.28 and 1.67, respectively (Figure 1R). Moreover, when Npr3-evoked activity was projected onto the sound-defined population dimensions, sound PC1 alone captured 82% of its reproducible variance, and the PC1 loading vectors were strongly aligned between sound and stimulation responses (median |cos θ| = 0.91; Figure 1R, Figure S4). Thus, natural and Npr3-evoked bursting occupy a shared low-dimensional population space, with burst amplitude providing the principal graded difference across inputs.

### Unilateral Npr3 stimulation recruits simultaneous bilateral BF bursting through recurrent circuit engagement

Having established that Npr3 stimulation accesses the endogenous structured BF burst mode, we next asked whether activation of only a subset of the circuit could nevertheless recruit the full structured burst. This provides a key test distinguishing feedforward and dynamical attractor accounts: feedforward recruitment requires effective inputs to access essentially the full bursting ensemble (Figure 1B), whereas recurrent circuit dynamics can allow activation of a restricted subset to recruit the full ensemble into the same structured burst (Figure 1C). Because Npr3 neurons showed no detectable projection to the contralateral BF (Figure 1D), unilateral Npr3 stimulation provided a spatially restricted perturbation that directly accessed only one hemisphere. We therefore asked whether unilateral stimulation would recruit bursting only within the directly accessed hemisphere, as expected from a feedforward account, or instead recruit the structured BF burst bilaterally through recurrent circuit engagement (Figure 2A).

**Figure 2.**
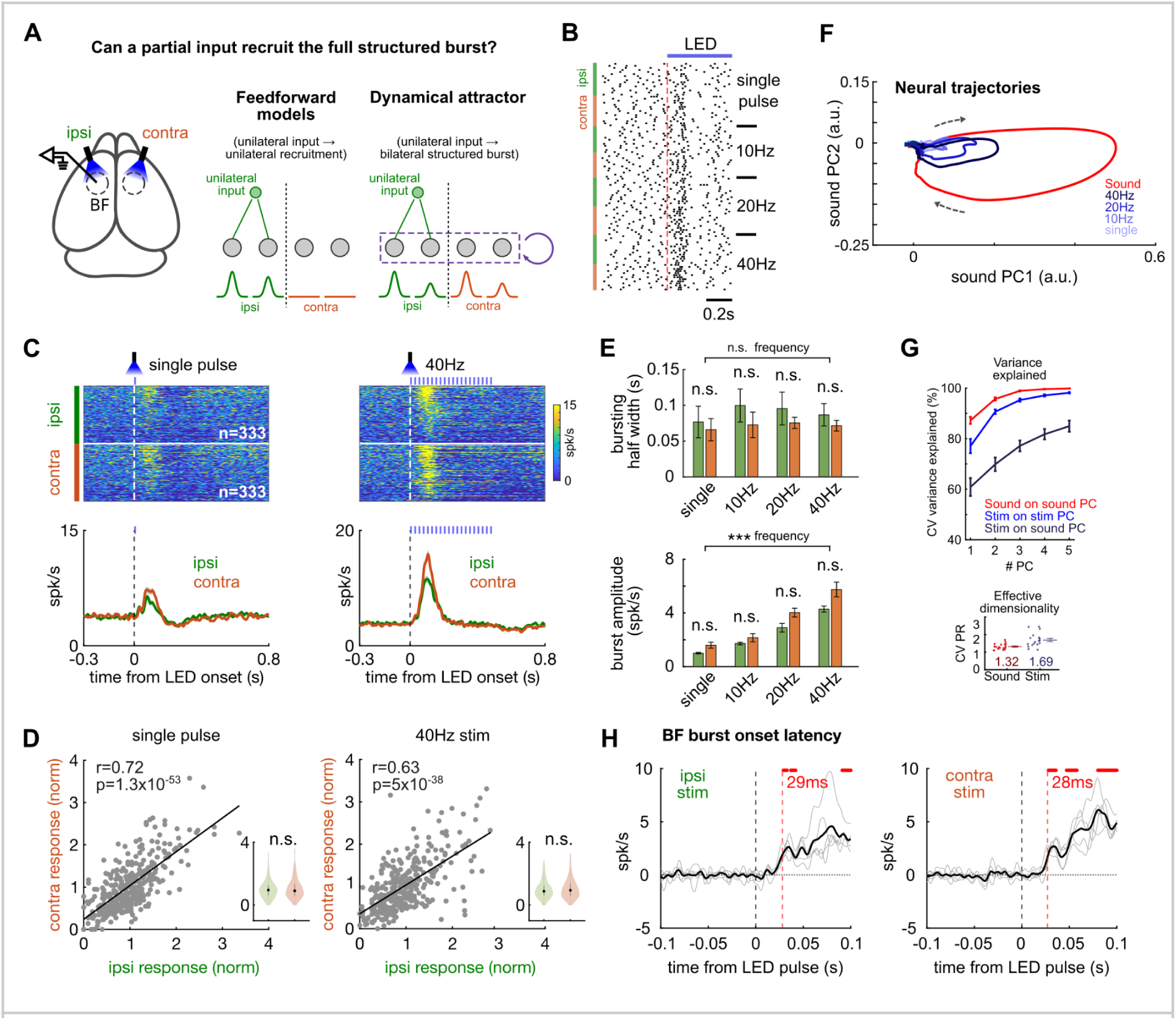
Unilateral Npr3 stimulation recruits bilateral BF bursting with matched onset latencies. **A,** Experimental design and model predictions for unilateral Npr3 stimulation. Npr3 neurons were stimulated ipsilateral or contralateral to the BF recording site. Feedforward models predict recruitment confined to the directly accessed hemisphere, whereas a dynamical attractor allows a unilateral input to recruit the full bilateral ensemble into the same structured burst through recurrent dynamics. **B,** Example BF bursting neuron showing similar responses to Npr3 stimulation delivered to the ipsilateral or contralateral BF. **C,** Population heatmaps and average PSTHs (mean±s.e.m.) showing BF bursting neuron responses to unilateral single-pulse and 40-Hz Npr3 stimulation. For each stimulation condition, responses of the same BF bursting neurons are shown for ipsilateral and contralateral stimulation, with neurons sorted identically. Unilateral stimulation recruited robust bursting in both hemispheres. **D**, Conserved cell-to-cell amplitude relationships between ipsilateral and contralateral stimulation responses, shown separately for single-pulse and 40 Hz Npr3 stimulation. Scatter plots show Pearson correlations between normalized response amplitudes evoked by ipsilateral and contralateral stimulation across individual BF bursting neurons. Insets compare paired ipsilateral and contralateral normalized response amplitudes across individual BF bursting neurons; no significant side bias was detected (Wilcoxon signed-rank test, p > 0.05; n = 333 neurons). **E,** Response half-width and mean firing rate of evoked BF bursting responses. Response half-width did not differ across stimulation frequencies or between stimulation sides (two-way repeated-measures ANOVA: frequency, *F*(3,12) = 0.43, *p* = 0.62; side, *F*(1,4) = 4.59, *p* = 0.099; frequency × side interaction, *F*(3,12) = 0.16, *p* = 0.82). Mean firing rate increased with stimulation frequency, with no significant difference between ipsilateral and contralateral stimulation (two-way repeated-measures ANOVA: frequency, *F*(3,12) = 160.57, *p* < 0.001; side, *F*(1,4) = 3.71, *p* = 0.13; frequency × side interaction, *F*(3,12) = 3.37, *p* = 0.10). **F,** Population neural trajectories, as in Figure 1Q, for mean sound responses (red) and unilateral Npr3 stimulation responses (blue). **G,** Variance explained and effective dimensionality, as in Figure 1R, for sound and unilateral stimulation responses. Mean ± SEM across 18 sessions; dots, sessions; group means shown below. **H,** Estimated onset latencies of population BF bursting responses to ipsilateral and contralateral stimulation. Gray lines represent individual animals (n=5), averaged across contributing stimulation protocols (see Figure S5 for details). Ipsilateral and contralateral onset latencies were closely matched, with both occurring after delays exceeding 25 ms.

Regardless of whether Npr3 stimulation was delivered ipsilateral or contralateral to the recording site (Figure 2A), both conditions evoked similar bursting responses in an example BF bursting neuron (Figure 2B) as well as across the population (Figure 2C). To compare cell-to-cell amplitude relationships, we normalized responses within each stimulation condition by the population burst amplitude of all simultaneously recorded BF bursting neurons across both hemispheres, thereby accounting for differences in stimulation efficacy due to opsin expression or fiber placement. Ipsilateral and contralateral stimulation preserved similar cell-to-cell amplitude relationships, with no detectable side-specific reorganization of the BF burst ensemble (Wilcoxon signed-rank test, p > 0.05; Figure 2D). Even without normalization, response half-width showed no significant effect of stimulation frequency or side, whereas mean firing rate increased with stimulation frequency but did not differ between ipsilateral and contralateral stimulation; neither measure showed a significant interaction between frequency and stimulation side (two-way repeated-measures ANOVA; Figure 2E). At the population level, unilateral Npr3 stimulation evoked trajectories closely resembling those of natural sound-evoked bursting, whether stimulation was ipsilateral or contralateral to the recording site (Figure 2F; Figure S4A). Cross-validated PCA confirmed that these responses remained strongly low-dimensional and closely aligned with the sound-defined population dimensions (Figure 2G; Figure S4B-D). Thus, unilateral Npr3 stimulation recruited the same bilaterally organized structured BF burst from either hemisphere.

The precise timing of Npr3 stimulation further revealed that BF burst onset was delayed and nearly identical for ipsilateral and contralateral responses (29 ms and 28 ms, respectively; Figure 2H; Figure S5), substantially slower than expected for immediate local recruitment. Together with the absence of detectable contralateral BF projections from Npr3 neurons (Figure 1D), this delayed and matched timing argues against a locally generated ipsilateral response that subsequently propagates to the opposite hemisphere, and instead implicates external circuit nodes beyond the BF. The matched ipsilateral and contralateral latencies further suggest that both ipsilateral and contralateral recruitment are likely mediated by a common long-range recurrent loop that extends from the BF through external circuit nodes and returns to both BF hemispheres. Such recurrent interactions provide a key circuit feature expected of a dynamical attractor.

### Spatially distinct Npr3 inputs are pooled into a common scalar BF output through approximate linear summation

Having established that the stable internal structure of the BF burst is generated by recurrent circuit dynamics of a dynamical attractor, we next asked how its flexible population burst amplitude is determined by input features. We had already shown that burst amplitude scaled with stimulation frequency at a single unilateral input site (Figure 2E). We therefore asked how inputs from spatially distinct sites combine when activated concurrently. Three possibilities were considered (Figure 3A): concurrent inputs could combine sublinearly through competition, supralinearly through synergy, or approximately linearly through independent contributions. These alternatives can be distinguished by comparing the observed concurrent response with the linear prediction obtained by summing the responses evoked by each input alone.

**Figure 3.**
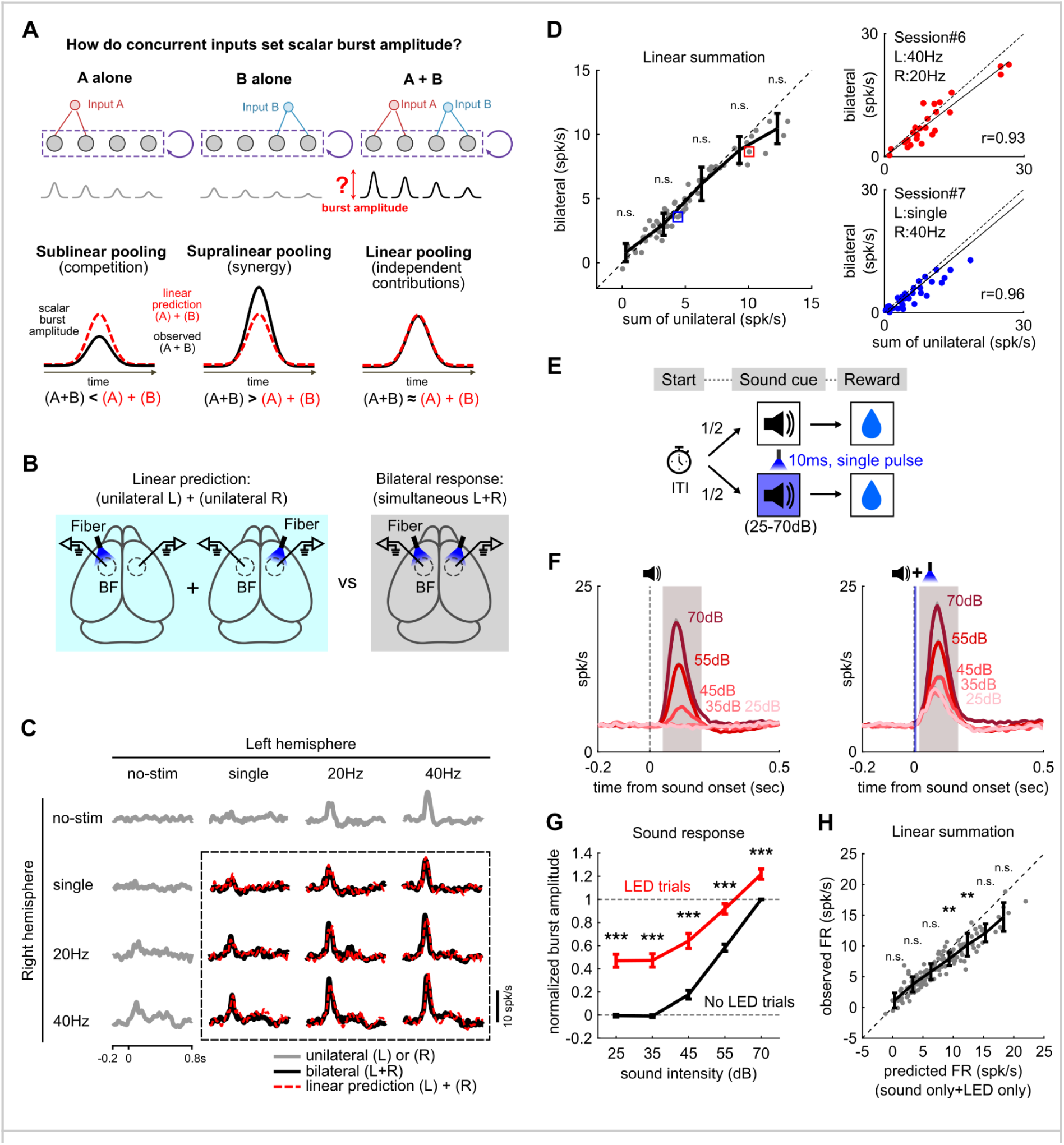
Concurrent inputs are pooled into a common scalar BF output through approximate linear summation. **A,** Possible rules by which concurrent inputs determine scalar BF burst amplitude. Inputs A and B may combine sublinearly (competition), supralinearly (synergy), or approximately linearly (independent contributions), defined by comparing the observed concurrent response (A+B; black) with the linear prediction from the summed individual responses, (A) + (B) (red dashed). **B,** Experimental test of spatial summation. The linear prediction, obtained by summing responses to left and right unilateral Npr3 stimulation, was compared with the empirically measured response to simultaneous bilateral stimulation. **C,** Population PSTHs from an example session in which the left and right BF hemispheres were stimulated independently at four levels: no stimulation, single pulse, 20 Hz, and 40 Hz, yielding 16 combinations. Gray traces show unilateral or no-stimulation responses; black traces within the dashed rectangle show bilateral responses; red dashed traces show the corresponding linear predictions obtained by summing the unilateral responses. **D,** Comparison of measured bilateral responses with summed-unilateral predictions. Left, summary across sessions and stimulation combinations. Gray dots represent individual session-condition observations from 7 sessions, each contributing nine bilateral stimulation conditions (63 observations total); black symbols show binned mean ± SEM. Binned observed-minus-predicted differences were tested against zero (Bonferroni-corrected Wilcoxon signed-rank tests, *p* > 0.05; *n* = 4–25 observations per bin). Right, neuron-level comparisons for two example stimulation conditions corresponding to the boxed observations at left, showing close agreement between measured and predicted responses. Dashed lines indicate unity. **E,** Auditory detection task testing summation between natural sensory and optogenetic inputs. A single 10-ms Npr3 stimulation pulse was delivered at sound onset on half of the trials. **F,** Population PSTHs (mean ± SEM) of BF bursting neurons evoked by sounds of different intensities on sound-only trials (left) and sound+LED trials (right). Shading indicates the window used to quantify burst amplitude. **G,** Normalized BF burst amplitude across sound intensities with (red) and without (black) Npr3 stimulation. Responses were normalized with the 70-dB sound response on no-LED trials set to 100% and baseline firing rate to 0% (Bonferroni-corrected Wilcoxon signed-rank tests; *n* = 36 sessions; \*\*\**p* < 0.001). **H,** Test of linear summation between natural sound and Npr3 inputs. Observed sound+LED responses were compared with predictions obtained by summing the separately measured sound-only and LED-only responses. Gray dots represent individual session-condition observations; black symbols show binned mean ± SEM. Responses followed the unity line over most of the operating range but became sublinear at higher predicted firing rates. Binned observed-minus-predicted differences were tested as in **D** (Bonferroni-corrected Wilcoxon signed-rank tests; *n* = 8–36 observations per bin; \*\**p* < 0.01; n.s., *p* > 0.05).

We tested these possibilities using spatially distinct Npr3 inputs from the two hemispheres (Figure 3B). Stimulation protocols were independently randomized between hemispheres, allowing unilateral and simultaneous bilateral responses to be measured within the same recording session (Figure 3C). We then compared the observed bilateral responses with the corresponding summed-unilateral predictions. At both the population and single-neuron levels, bilateral Npr3 responses were well approximated by the summed unilateral prediction across a wide dynamic range (Bonferroni-corrected Wilcoxon signed-rank tests, p > 0.05; Figure 3D; Figure S6). Thus, spatially distinct Npr3 inputs did not compete or interact synergistically, but instead combined through approximate linear summation when activated concurrently.

### Natural and optogenetic inputs are pooled by approximate linear summation during behavior

Having shown that two spatially distinct Npr3 inputs combine by approximate linear summation outside the behavioral task, we next asked whether the same integration rule extends to a natural reward-predicting stimulus during behavior. We trained head-fixed mice to perform an auditory detection task in which sounds of varying intensity predicted the same reward, providing a graded natural input. On half of the trials, a single 10-ms Npr3 stimulation pulse was delivered at sound onset, allowing direct comparison of sound-only and sound+Npr3 responses across sound intensities (Figure 3E).

Sound-evoked BF burst amplitude increased systematically with sound intensity (Figure 3F), consistent with previous findings^4^. Adding a single Npr3 pulse increased burst amplitude across sound intensities while preserving the overall temporal profile of the response (Bonferroni-corrected Wilcoxon signed-rank tests, all p < 0.001; Figure 3F,G). We then asked whether the combined response could be predicted by linear summation of the two inputs. Because 25- and 35-dB sounds did not elevate BF activity above baseline, sound+Npr3 responses at these intensities were used to estimate the Npr3-only component. Adding this component to the corresponding sound-only responses closely predicted the observed sound+Npr3 responses across the main operating range (Figure 3H). The relationship became sublinear as population burst amplitude approached saturation, consistent with a physiological ceiling, and the range of approximate linearity matched that observed during bilateral Npr3 stimulation (Figure 3D). Thus, approximate linear summation extended from spatially distinct Npr3 inputs to combinations of natural and optogenetic inputs during behavior, revealing an integration principle used by endogenous reward-predicting inputs within the BF burst mode.

### Npr3-induced BF bursting enhances reward prediction during behavior

Having shown that Npr3 stimulation recruits the endogenous BF burst mode and increases its scalar amplitude during behavior, we next asked whether Npr3-induced bursting conveys the same reward-prediction information as natural BF bursting (Figure 4A). In the auditory detection task, higher sound intensities support stronger reward prediction and evoke larger BF bursts, which are associated with higher hit rates, shorter reaction times, and smaller subsequent reward responses^4^. If Npr3-induced bursting engages the same reward-prediction signal, increasing BF burst amplitude on stimulated trials should shift behavioral performance and subsequent reward responses in the same directions.

**Figure 4.**
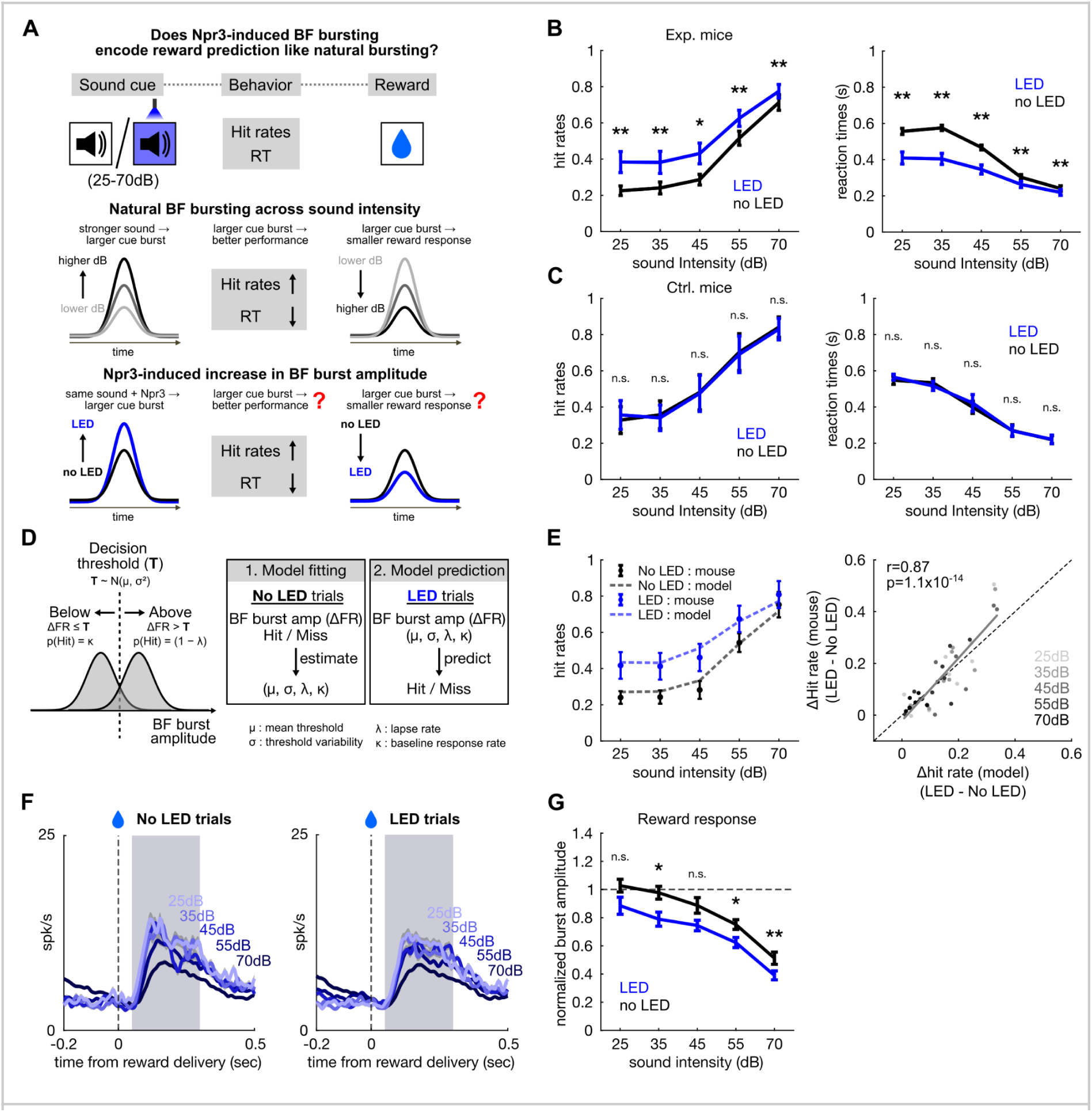
Npr3 activation improves behavioral performance by enhancing reward prediction. **A,** Test of whether Npr3-induced BF bursting encodes reward prediction like natural BF bursting. Across sound intensities, larger natural cue bursts are associated with higher hit rates, shorter reaction times, and smaller subsequent reward responses. If Npr3-induced bursting engages the same reward-prediction signal, it should similarly improve behavioral performance and reduce the subsequent reward response. **B,** Behavioral effects of Npr3 stimulation. Npr3 activation increased hit rate (left) and shortened reaction time (right) across sound intensities relative to no-LED trials (Bonferroni-corrected Wilcoxon signed-rank tests; *n* = 10 animals; \**p* < 0.05; \*\**p* < 0.01). **C,** Same analysis in control mice expressing EYFP without opsin. LED stimulation did not significantly alter hit rate or reaction time across sound intensities (Bonferroni-corrected Wilcoxon signed-rank tests; *n* = 6 animals; n.s., *p* > 0.05). **D,** Model-fitting and prediction strategy. Behavioral responses were modeled using a probabilistic decision threshold, with trial-by-trial BF burst amplitude compared with a latent threshold T∼N(μ,σ2). Model parameters, including threshold mean and variability, lapse rate, and baseline response rate, were estimated from no-LED trials and then applied without refitting to LED trials to predict behavioral responses from the observed BF burst amplitudes. **E,** Model performance. Left, observed hit rates and model predictions across sound intensities for no-LED and LED trials. Right, comparison of the observed and predicted LED-induced changes in hit rate across animals and sound conditions (*n* = 9 animals); *r*, Pearson correlation coefficient. **F,** Reward-aligned population PSTHs (mean ± SEM) of BF bursting neurons on no-LED (left) and LED (right) trials across sound intensities. Shading indicates the window used to quantify reward-response amplitude. **G,** Normalized reward-response amplitude across sound intensities with (blue) and without (black) Npr3 stimulation. Responses were normalized with the reward response on 25- and 35-dB no-LED trials set to 100% and baseline firing rate to 0%. Npr3 stimulation reduced subsequent reward responses, consistent with stronger reward prediction producing a smaller reward prediction error (Bonferroni-corrected Wilcoxon signed-rank tests; *n* = 36 sessions; \**p* < 0.05; \*\**p* < 0.01; n.s., *p* > 0.05).

Consistent with this prediction, a single 10-ms Npr3 stimulation pulse at sound onset increased hit rate and reduced reaction time across sound intensities in experimental mice (Bonferroni-corrected Wilcoxon signed-rank tests, all p < 0.05; Figure 4B), whereas the same manipulation produced no detectable effect in control mice (Figure 4C). Additional control analyses argued against simple sensory or motor accounts (Figure S7). Npr3 stimulation increased responding even on 0-dB trials, in which no auditory stimulus was present, arguing against enhanced sensory gain.

Conversely, stimulation-associated licking probability was substantially lower outside the behavioral task than during stimulated trials in the task, arguing against a simple motor explanation. Thus, a single 10-ms Npr3 stimulation pulse robustly improved behavioral performance, consistent with enhanced reward prediction.

If this behavioral improvement reflected stronger reward prediction encoded by the Npr3-induced increase in cue-period BF bursting, then the accompanying increase in burst amplitude should quantitatively predict the change in behavior. Motivated by the observation that cue-period BF responses were larger on hit than miss trials^4^ (Figure S8), we used a probabilistic decision-threshold model to link trial-by-trial BF burst amplitude to behavioral outcome, in which a reward-seeking response was generated when burst amplitude exceeded a latent threshold (Figure 4D; Figure S8). For each session, model parameters were estimated solely from non-stimulated sound-only trials based on the relationship between population burst amplitude and behavioral outcome, and then applied without refitting to Npr3-stimulated trials from the same session. The model not only reproduced hit rates in sound-only trials but also accurately predicted the Npr3-induced increase in hit rate across sound intensities, with predicted and observed stimulation effects closely matched across conditions (Figure 4E). It also reproduced how cue-period BF activity varied with behavioral outcome and sound intensity in both sound-only and stimulated trials (Figure S8). Thus, the behavioral improvement produced by Npr3 stimulation was quantitatively accounted for by the accompanying increase in BF burst amplitude, supporting the conclusion that Npr3-induced BF bursting encodes reward prediction like natural BF bursting.

A final test came from reward-period responses. If cue-related reward prediction is stronger, the BF bursting response to the subsequent reward should be reduced, an inverse relationship characteristic of reward-prediction-error encoding and previously demonstrated in BF bursting neurons^4–7^. In non-stimulated sound-only trials, reward-period bursting declined systematically with increasing sound intensity, consistent with stronger reward prediction for higher-intensity cues (Figure 4F,G). In Npr3-stimulated trials, reward-period responses were further reduced across all sound conditions (Figure 4F,G), even though stimulation was delivered only transiently at sound onset and not at reward delivery. Thus, both natural increases in cue strength and optogenetic augmentation of cue-evoked BF bursting produced the same reduction in subsequent reward responses, consistent with stronger reward prediction assigned to the cue. Together with the behavioral modeling above, these findings identify population burst amplitude as the scalar output through which the shared BF burst mode encodes reward-prediction strength, demonstrating that Npr3 stimulation enhances endogenous reward prediction by engaging this shared mode.

## Discussion

In this study, we asked how BF bursting reconciles two competing constraints: stable internal structure and flexible population burst amplitude. We tested the hypothesis that this organization is generated by an internally organized dynamical attractor rather than imposed directly by feedforward inputs. Using BF Npr3 neurons as a defined experimental entry point, we found that Npr3 stimulation recruited the endogenous BF burst mode, preserving cell-to-cell amplitude relationships, temporal envelope, and low-dimensional population dynamics while allowing burst amplitude to scale with input strength (Figure 1). Unilateral stimulation recruited the structured burst bilaterally with matched onset latencies, implicating recurrent interactions beyond the BF (Figure 2), while concurrent optogenetic and natural inputs summed approximately linearly in burst amplitude (Figure 3). During behavior, Npr3-induced increases in burst amplitude enhanced reward prediction and behavioral performance (Figure 4). Together, these findings demonstrate that a dynamical attractor generates BF bursting, combining an internally constrained burst structure with a flexible scalar amplitude that integrates concurrent inputs and encodes reward-prediction strength.

### BF ensemble bursting as an internally organized dynamical attractor

Attractor-dynamics frameworks emphasize that internally organized circuit states can converge across distinct inputs onto a shared population trajectory, evolve along constrained temporal envelopes, respond in a state-dependent manner once engaged, and often depend on recurrent circuit interactions for their recruitment^8–10,29^. Several features of BF bursting fit this framework. First, distinct natural and optogenetic inputs converged onto the same low-dimensional burst trajectory, preserving its temporal envelope and cell-to-cell amplitude relationships despite differences in input source and pattern (Figures 1–3). Second, once this trajectory was engaged, additional Npr3 pulses had progressively reduced effects (Figure 1), revealing the nonlinear state dependence expected when subsequent inputs act on an evolving circuit state rather than on a passive linear filter. Third, unilateral Npr3 stimulation recruited bilateral bursting with delayed and matched ipsilateral and contralateral onset latencies, implicating recurrent circuit interactions beyond the directly stimulated BF (Figure 2).

Critically, these findings are difficult to reconcile with feedforward models in which the output pattern is imposed by the initiating input. Unilateral Npr3 stimulation provided a direct test of whether an effective input must access the full bursting ensemble. Because Npr3 neurons showed no detectable projection to the contralateral BF, unilateral stimulation directly accessed only one hemisphere, yet nevertheless recruited the same structured burst bilaterally (Figure 2). Thus, an input need not directly reach the full ensemble to recruit the full structured burst. Together, these properties indicate that the defining structure of the BF burst is generated by circuit dynamics rather than specified by the initiating input. A dynamical attractor therefore allows diverse inputs to recruit a shared BF population response. This convergence provides a natural explanation for how reward-predicting stimuli from different sensory modalities can recruit the same BF bursting ensemble in a modality-invariant manner^7^.

The attractor underlying BF bursting shares important features with other attractor systems that integrate inputs along a dominant low-dimensional axis, while differing in potentially important aspects of its dynamics. Examples include line attractors such as the brainstem oculomotor integrator and the aggressive-state attractor in the ventromedial hypothalamus^11,30,31^, as well as prefrontal decision circuits that accumulate sensory evidence along a slow dimension between choice attractors^32–34^. However, the BF burst differs from these systems in two important ways. First, BF bursting is a transient event lasting a few hundred milliseconds rather than a slow integrator operating over seconds to minutes. Second, its integration window is self-terminating: once the burst trajectory is engaged, later inputs lose efficacy. Whether BF bursting represents a transient variant of this attractor family or a distinct dynamical regime remains to be determined.

### Linear summation of population burst amplitude supports source-invariant reward prediction

Although attractor dynamics are inherently nonlinear and state dependent, BF ensemble bursting nonetheless contained a surprisingly linear dimension in its flexible population burst amplitude. For a single input source, burst amplitude scaled with input strength, whether defined by Npr3 stimulation frequency (Figures 1–2) or sound intensity (Figure 3). When multiple sources were activated concurrently, their contributions summed approximately linearly along the burst-amplitude dimension, whether they arose from spatially distinct Npr3 populations or from Npr3 stimulation combined with a natural reward-predicting sound, becoming sublinear only as responses approached a physiological ceiling (Figure 3H). Importantly, the latter experiment showed that the same integration rule operates during endogenous reward prediction, rather than being restricted to optogenetically imposed inputs. Consistent with this principle, auditory and visual inputs also combine approximately linearly in BF bursting during behavior^7^.

This linear summation reveals a simple algorithm for computing reward-prediction magnitude. Natural sensory inputs are likely represented across distributed, relatively sparse neuronal populations, so individual input populations may have access to only part of the BF circuit. Their contributions can nevertheless be pooled approximately linearly within the attractor into a graded measure of predictive strength, while recurrent dynamics preserve the shared temporal and cell-to-cell structure of the burst. In this sense, linear summation provides an arithmetic operation for constructing reward prediction from distributed inputs. This complements the influential finding that dopamine reward-prediction errors can be generated through subtraction of expected from received value^35^: whereas subtraction provides an operation for computing prediction error, our results identify summation as an operation for constructing reward-prediction magnitude. Approximate linear summation may therefore allow the BF to transform heterogeneous predictive inputs into a common, source-invariant scalar reward-prediction signal.

### Circuit mechanisms should be sought in reciprocal loops

The attractor interpretation shifts the mechanistic question from identifying a single proximal feedforward input that directly drives BF bursting to identifying the reciprocal circuit interactions that allow diverse inputs to recruit the same structured population mode. This view implies that the native BF burst may depend on recurrent interactions across multiple brain regions rather than circuitry contained within the BF itself. Isolated BF preparations may therefore be insufficient to reproduce the circuit conditions that generate the full population event. Major projection targets of Npr3 neurons provide a restricted set of candidate nodes for such reciprocal interactions, including frontal cortex, the thalamic reticular nucleus, and the midbrain ventral tegmental area (VTA). Determining which of these pathways participate in reciprocal interactions that support burst recruitment, and which primarily carry downstream consequences of the burst, will be important for defining the circuit architecture of the attractor.

The reciprocal-loop view also changes how the initiating events should be sought. Because Npr3-evoked BF bursting emerged only after delays exceeding 25 ms and could be triggered by a single brief stimulation pulse, the causally relevant circuit activity may begin well before the burst becomes detectable in the BF. Such activity could be sparse, distributed across several regions, transient, or small in amplitude, and therefore missed by analyses focused only on strong activity immediately preceding burst onset. Mechanistic studies should therefore search for earlier circuit events that enable entry into the BF burst state, rather than only for strong inputs arriving immediately before the population burst.

### Npr3 neurons provide an entry point into the BF burst attractor

Npr3 neurons provided a defined experimental entry point into the dynamical attractor underlying BF bursting, allowing its organization to be probed outside the behavioral events that normally evoke BF bursting. This was critical for controlling the timing, strength, spatial origin, and combination of inputs in ways that are difficult to achieve with natural stimuli. Npr3 stimulation therefore served primarily as a perturbation for revealing the circuit dynamics underlying BF bursting. Importantly, our results do not imply that Npr3 neurons constitute the only entry point into the attractor or are necessary for endogenous burst initiation. Rather, their value here is that selective activation of this population was sufficient to expose a dynamical regime in which restricted inputs recruit the same structured BF burst. Regardless of whether Npr3 neurons are required for natural bursting, our conclusion that BF population bursting is generated by a dynamical attractor remains unchanged.

We selected Npr3 from single-cell and spatial transcriptomic datasets^12–15,20,21^ because its distinctive spatial expression pattern follows the anatomical organization of the broader BF macrosystem across its entire anteroposterior extent (Figure S1), extending from the medial septum and diagonal band anteriorly, through the substantia innominata and magnocellular preoptic area, to the external globus pallidus posteriorly^16,22^. This distribution suggests that Npr3 marks a distinct, distributed GABAergic population associated with the broader BF macrosystem rather than a cell type confined to a single anatomically defined nucleus. Consistent with this interpretation, Npr3+ neurons in the external globus pallidus have been identified as a GABAergic projection population with similar outputs, including prominent innervation of the cortex, thalamic reticular nucleus and SNc^23,24^.

What remains unresolved is the role of Npr3 neurons during behavior and their relationship to the broader BF bursting population. Because phototagged Npr3 neurons were too sparse in our recordings for population-level characterization (Figure S3), we could not determine from electrophysiological recordings whether Npr3 neurons normally participate in the endogenous burst ensemble. One possibility is that they act primarily as an input population that recruits the bursting ensemble without constituting a major part of it. Alternatively, they may represent a subset of BF bursting neurons and therefore provide one of several access points into the shared population mode. Distinguishing these possibilities will require molecular markers that directly identify BF bursting neurons, which likely constitute a distinct subset of BF GABAergic neurons^18^.

## Materials and Methods

### Animals

A total of 26 Npr3-IRES2-Cre-D mice^36^ (Npr3-Cre, Jax #031333; 18 male and 8 female; 2-4 months old at the time of surgery) were used in this study. See Table S1 for animal contributions to each experiment and analysis. Mice were housed on a reversed 12-h light/dark cycle, and experiments were performed during the dark phase. Food was available ad libitum. Water was provided ad libitum except during behavioral training and recording procedures, when mice were water restricted to maintain 80% of their baseline body weight^37^. During water restriction, mice received 1 ml of water per day, including water obtained during the daily 1-h behavioral training session and supplemental water provided after training. All experimental procedures were approved by the Institutional Animal Care and Use Committee of National Yang Ming Chiao Tung University, Taiwan (NYCU IACUC #1080510, #1131207 and #1140404).

### Apparatus

Behavioral experiments were conducted in a head-fixed mouse setup adapted from previously described designs^37^. The setup was housed within light- and sound-attenuating cubicles measuring 55 × 43.5 × 54 cm (L × W × H). Each cubicle was equipped with an exhaust fan (Sanyo Denki America Inc., USA; Cat. No. 9S1224M401) that also served to mask external noise. Mice were head-fixed above a custom behavioral platform, with an infrared (IR) sensor positioned in front of the tongue to detect licking (IR emitter, ams OSRAM, Germany; Cat. No. SFH 4851; IR receiver, Vishay, USA; Cat. No. BPW76A). Water rewards were delivered through a lick spout positioned in front of the animal and controlled by a syringe pump^38^. The pump was calibrated to dispense 0.6 μl of water per motor step.

A speaker (Visaton, Germany; Cat. No. 2240) was positioned in front of the animal and controlled by a programmable audio generator (ANL-926; Med Associates, USA). A white light-emitting diode (LED) lamp (VCC, USA; Cat. No. VC1860245W3D) positioned near the speaker served as the visual stimulus. Behavioral training and task control were implemented using Med-PC software (Version V; Med Associates, USA), which recorded all event timestamps with 1-ms resolution and sent TTL signals to the electrophysiological recording and optogenetic stimulation systems.

### Stereotaxic surgery

Mice underwent stereotaxic surgery for viral injection and implantation of optic fibers or optrodes before behavioral training. Surgeries were performed under isoflurane anesthesia (4% for induction and 1-1.5% for maintenance). Eyes were protected with ophthalmic ointment during surgery. After shaving, the scalp was disinfected with 70% ethanol followed by 10% povidone-iodine. An incision was made to expose the skull, and the skull was leveled by aligning bregma and lambda to the same dorsoventral (DV) coordinate. Craniotomies were made above the bilateral basal forebrain target region centered on the SI (BF; AP: −0.1 mm, ML: ±1.5 mm relative to bregma) using a dental drill. Virus injections were performed using a programmable nanoliter injector (Nanoject III; Drummond Scientific, USA). The injection pipette was lowered at 10 μm/s to the target depth (DV: −4.7 mm from cortical surface). Virus was injected at 1 nl/s, and the pipette was left in place for 10 min after injection before being gradually withdrawn. Optic fibers and electrode implants were secured to the skull with Super-Bond (Sun Medical, Japan). A custom stainless-steel headbar was then attached using Super-Bond and dental cement for head-fixed behavioral experiments. All mice received ibuprofen for postoperative analgesia and were allowed to recover for at least 2 weeks before further experiments.

For anatomical tracing experiments, AAV5-EF1α-DIO-EYFP (Addgene, USA; #27056; titer ≥1 × 10^13 gc/ml) was injected unilaterally into the BF at a depth of DV−4.7 mm (50 nl), followed by a 4-week expression period (Figure 1D). For optogenetic manipulation experiments, AAV5-EF1α-DIO-hChR2(H134R)-EYFP (Addgene, USA; #20298; titer ≥1 × 10^13 gc/ml) or the control vector AAV5-EF1α-DIO-EYFP (Addgene, USA; #27056; titer ≥1 × 10^13 gc/ml) was injected bilaterally into the BF (100 nl per site). Optic fibers with ceramic ferrules (200-μm core diameter, 0.66 NA; Inper, China) were implanted above the BF (AP: −0.1 mm, ML: ±1.5 mm, DV: −4.4 mm).

For electrophysiology experiments, AAV5-EF1α-DIO-hChR2(H134R)-EYFP was injected into the bilateral BF (100 nl per site). The optrode assembly consisted of two bundles, each containing a 200-μm optic fiber surrounded by 14 polyimide-insulated tungsten wires (38 μm in diameter; California Fine Wire, USA). The electrodes and optic fiber were glued together and enclosed in 28-gauge polyimide tubing, with the electrode tips positioned 250 μm deeper than the fiber tip. Each optrode bundle was controlled by a precision microdrive, allowing gradual electrode advancement after surgery. Electrode impedances were ∼0.1 MΩ at 1 kHz, measured using an Open Ephys Acquisition Board^39^. Pin connectors (#1372-2; Keystone Electronics, USA) secured to the skull served as the electrical reference and ground: one connector positioned over the cerebellum served as the reference, and another positioned over the frontal cortex served as the ground. During implantation, the electrode bundles were retracted inside the cannulae, and the cannulae were lowered to an initial depth of 3 mm below the cortical surface using a micropositioner (Biochiefdom International Ltd., Taiwan), at 5 μm/s for the first 2 mm and 2 μm/s for the final 1 mm. Once the target depth was reached, the electrode assembly and skull screws were secured with Super-Bond. The electrodes and optic fibers were then advanced through the microdrives to a final depth of 4 mm below the cortical surface.

### Behavioral task

Mice were trained in three stages. In the first stage, mice were habituated to head fixation for 1-2 days and trained to lick the spout to obtain water rewards. Each water delivery was paired with a 6-kHz pure tone cue (0.5 s, 80 dB), and each reward consisted of 2.4 μl of water.

In the second stage, mice were trained on a sound-reward association task for 5-7 days. Each trial consisted of a 6-kHz pure tone cue (0.5 s, 70 dB) followed by a 3-s response window. Three consecutive licks within the response window triggered water delivery, with reward delivered upon the third lick. Trials with licking responses terminated 1 s after the last lick, whereas trials without licking responses terminated at the end of the 3-s response window. Each trial was followed by an unsignaled inter-trial interval (ITI) of 3-5 s, and licking during the ITI reset the ITI timer. Mice advanced to the final auditory detection task after reaching a criterion of 100 hit trials per day for 2 consecutive days.

In the third stage, mice performed the auditory detection task. Each trial consisted of a 6-kHz pure tone cue (0.5 s) presented at one of five sound intensities: 25, 35, 45, 55, or 70 dB. To maintain stable behavioral performance across sessions, sound intensities were presented with unequal probabilities: 70 and 55 dB trials each occurred with a probability of 1/3, whereas 45, 35, and 25 dB trials each occurred with a probability of 1/9. Three consecutive licks within a 1.5-s response window after cue onset triggered water delivery, with reward delivered upon the third lick. Reward volume was the same across all sound intensities, with each rewarded trial delivering 2.4 μl of water. Trials with licking responses terminated 1 s after the last lick, whereas trials without licking responses terminated at the end of the 1.5-s response window. Each trial was followed by an unsignaled ITI of 3-5 s, and licking during the ITI reset the ITI timer. In a subset of experiments, the 25 dB condition was replaced with a 0 dB condition, corresponding to the absence of an auditory cue, to assess behavior in the absence of sensory input (Figure S7).

### Optogenetic stimulation

For the stimulation-only protocol, Npr3 neurons were optogenetically stimulated after completion of the behavioral task, in the absence of auditory cues or rewards (Figures 1 and 2). Blue light (465 nm, 4–6 mW) was delivered using a PlexBright LED system (Plexon Inc., USA). Four Npr3 stimulation conditions were used: a single 10-ms pulse or 500-ms pulse trains at 10, 20, or 40 Hz, with a pulse duration of 10 ms (Figure 1N). For the experiments in Figure 1, stimulation was delivered bilaterally. For the experiments in Figure 2, the same stimulation protocols were delivered unilaterally to either the left or right BF. Conditions were randomly interleaved with an intertrial interval (ITI) of 3–5 s, and each condition was repeated for 15 trials.

To assess approximate linear summation between spatially distinct Npr3 inputs, an additional stimulation-only experiment was performed in which the left and right BF hemispheres were stimulated independently (Figure 3B,C; Figure S6). Four stimulation conditions were applied independently to each hemisphere: no stimulation, single pulse, 20 Hz, or 40 Hz, yielding 16 possible bilateral combinations. The 16 conditions were presented in random order.

For Npr3 stimulation during the auditory detection task, a single 10-ms stimulation pulse was delivered at sound onset on a randomly selected 50% of trials (Figure 3E). The remaining trials were sound-only trials without optogenetic stimulation.

### Electrophysiological recording

Extracellular signals were recorded from the BF, band-pass filtered from 0.3 Hz to 7.5 kHz, amplified using an Intan headstage (#C3324, Intan Technologies, USA), and digitized at 30 kHz with the Open Ephys Acquisition Board^39^. Spike sorting was performed offline using the KlustaKwik algorithm^40^, followed by manual curation in a custom Python-based graphical user interface. Single units were identified based on clear separation from noise clusters and minimal refractory period violations (<0.1% of spikes occurring within a 1.5-ms interspike interval). Cross-correlation analyses were performed to eliminate duplicate units recorded simultaneously across multiple electrodes. Units with mean firing rates below 0.1 spikes/s were excluded from further analysis.

### Perfusion and histology

At the end of the experiment, mice were deeply anesthetized with an overdose of sodium pentobarbital (150 mg/kg, i.p.) and transcardially perfused with phosphate-buffered saline (PBS), followed by 4% paraformaldehyde (PFA). Brains were removed, postfixed in 4% PFA overnight at 4 °C, and then cryoprotected sequentially in 20% sucrose in PBS for 1 day and 30% sucrose in PBS for 2 days. Brains were frozen and coronally sectioned at 50 μm using a sliding microtome (SM2010 R; Leica Biosystems, Germany).

Free-floating sections were washed in PBS containing 0.2% Triton X-100 and blocked for 1 h at room temperature in PBS containing 5% normal donkey serum (Jackson ImmunoResearch, USA; Cat. No. 017-000-121; RRID: AB_2337258). Sections were then incubated overnight at 4 °C with a chicken anti-GFP primary antibody (Abcam, UK; Cat. No. ab13970; RRID: AB_300798; 1:1,000). Sections were rinsed three times in PBS for 10 min each and incubated for 2 h at room temperature with an Alexa Fluor 488-conjugated donkey anti-chicken secondary antibody (Thermo Fisher Scientific, USA; Cat. No. A78948; RRID: AB_2921070; 1:1,000). Sections were then washed, mounted with Vectashield antifade mounting medium (Vector Laboratories, USA; Cat. No. H-1200-10), and coverslipped.

Viral expression and the locations of optic fibers and optrode recording sites were verified histologically using fluorescence microscopy (Axio Imager M2, Carl Zeiss Microscopy, Germany; M205 FCA, Leica Microsystems, Germany).

### Data analysis

Data were analyzed using custom MATLAB scripts (R2019b; The MathWorks Inc., USA).

#### Session selection criteria

For electrophysiological recordings, only sessions with electrode positions separated by at least 125 μm were included in the analysis, to minimize repeated sampling of the same neuronal populations across sessions. For analyses of population dimensionality and subspace alignment, only sessions containing at least five BF bursting neurons were included, resulting in the exclusion of one session from the unilateral stimulation response analysis. For behavioral experiments, sessions with fewer than 80 hit trials were excluded from analysis. This criterion excluded 7 sessions, including 5 sessions from the 25–70 dB task and 2 sessions from the 0–70 dB task. The numbers of animals and sessions included in each experiment are summarized in Table S1.

#### Identification of BF bursting neurons

BF bursting neurons were defined as BF single units exhibiting a cue-evoked increase in firing rate of more than 2 spikes/s during the [0.05, 0.2]-s window after cue onset, compared with baseline firing rate measured during the [−2, −1]-s window relative to all stimulus onsets. In addition, BF bursting neurons were required to have a baseline firing rate between 0.1 and 10 spikes/s, based on previously established criteria^4–7,25,26^.

#### BF response amplitudes and their normalization

BF bursting responses were quantified as the mean firing rate within predefined post-event windows. Sound-evoked responses were measured during the [0.05, 0.2]-s window after sound onset, reward responses during the [0.05, 0.3]-s window after reward delivery, and Npr3-evoked responses during the [0.02, 0.17]-s window after stimulation onset.

To compare cell-to-cell amplitude relationships across conditions (Figures 1L,M and 2D), each BF bursting neuron’s response amplitude was normalized by the mean response amplitude of all simultaneously recorded BF bursting neurons across both hemispheres in the same condition. This normalization was applied separately to sound-evoked and Npr3-evoked responses.

For analyses of population BF burst amplitude, response amplitude was calculated within each session by averaging firing rates across BF bursting neurons and subtracting the corresponding baseline firing rate (Figures 1P, 2E, 3D, 3H and 4D). For analyses requiring comparison across sessions during the auditory detection task, these baseline-subtracted response amplitudes were further normalized to a reference response within the same session. Sound responses at each sound intensity were normalized to the mean burst amplitude in 70-dB sound-only trials (Figure 3G), whereas reward responses were normalized to the mean reward response amplitude in low-intensity (25- and 35-dB) sound-only trials (Figure 4G).

Burst half-width (Figure 1O) was defined as the temporal width of the response measured at half-maximum amplitude. Peak firing rate was defined as the maximum firing rate during the [0.02, 0.17]-s window after stimulation onset. Half-maximum was calculated as the firing rate halfway between baseline and peak firing rate.

#### Population dimensionality and subspace alignment

To visualize population trajectories evoked by natural sound and Npr3 stimulation (Figures 1Q, 2F and S4A), spike trains were binned and trial-averaged separately for each stimulus condition within each session to obtain peristimulus time histograms (PSTHs; time × neurons), which were smoothed with a Hann window (40 bins) along time. Neurons were then pooled across sessions into a pseudo-population, and each condition was projected onto the leading two principal components (PCs) of the sound-evoked pseudo-population response. This pseudo-population analysis was used only for trajectory visualization; all quantitative analyses of dimensionality and subspace alignment described below were performed separately within each session using simultaneously recorded neuronal populations.

To quantify population dimensionality and determine how much Npr3-evoked activity occupied the dimensions defined by natural sound-evoked bursting (Figures 1R, 2G and S4B,C), we used cross-validated PCA (cvPCA)^27^ within each recording session. For each condition, single trials were randomly partitioned into two disjoint halves (50/50 split, 200 repetitions). PCs were defined from the trial-averaged covariance of one half, and the cross-validated eigenvalue of each PC was computed as the covariance between the two halves’ projections onto that PC. Because sampling noise is independent between halves, its expected contribution to this cross-covariance is zero, thereby isolating reproducible population variance. To compare conditions on an equal footing, trial counts were subsampled to the minimum shared by the compared conditions within each session before splitting. Cross-validated cumulative variance explained was calculated as the running sum of the leading cross-validated eigenvalues, normalized by the total cross-validated variance, and evaluated for the first five PCs (k = 1–5). Effective dimensionality was summarized as the cross-validated participation ratio (CV PR), PR = (Σλ)²/Σλ², where λ denotes the cross-validated eigenvalue of each principal component, reflecting its reproducible variance contribution, computed over the full-rank cross-validated eigenvalue spectrum^28^.

The same cross-validated framework was used to quantify how much reproducible stimulation variance was captured by sound-defined population dimensions. For each repetition, stimulation responses from the two trial halves were projected onto (i) PCs defined from one half of the stimulation trials, providing the condition-specific ceiling, and (ii) PCs defined from the complete set of sound trials, providing a measure of alignment with the sound-defined population dimensions. Cross-validated stimulation variance was calculated as the covariance between the two stimulation trial halves after projection onto each set of PCs. Both estimates were normalized by the same total cross-validated stimulation variance. Because the same stimulation trial halves and normalization were used for the ceiling and alignment estimates, residual trial noise contributed equivalently to both and was approximately zero in expectation. The ratio of variance captured by sound-defined versus stimulation-defined dimensions therefore provides a measure of subspace alignment; for a single dimension, this ratio approximates cos²θ between the corresponding loading vectors. Values were averaged first across cross-validation repetitions and then across sessions for population-level summaries.

To directly quantify alignment of the dominant population pattern between sound- and stimulation-evoked activity (Figures 1R and S4D), we calculated the absolute cosine similarity, |cos θ|, between their leading PC1 loading vectors within each session. Significance was assessed against a null distribution generated by randomly permuting neuron identity in the stimulation PC1 loading vector (1,000 permutations). A session-specific upper bound, representing the expected ceiling imposed by finite trial sampling, was obtained by calculating the same cosine similarity between two independent, trial-count-matched split-halves of the sound response. PC1 alignment was quantified separately for bilateral, ipsilateral and contralateral Npr3 stimulation.

#### BF burst onset latency following Npr3 stimulation

To estimate BF burst onset latency following an Npr3 stimulation pulse (Figure S5), we combined the original stimulation-evoked responses with contrast-derived responses that isolated responses to later pulses within stimulation trains. For each session and stimulation condition, BF bursting neuron population PSTHs were first baseline subtracted. Four original responses were obtained from the first stimulation pulse in each of the four stimulation protocols: single pulse, 10 Hz, 20 Hz, and 40 Hz.

To generate additional latency estimates, we computed pairwise differences between stimulation protocols that shared earlier pulse timing but differed at a later pulse: 10 Hz minus single pulse, 20 Hz minus single pulse, 40 Hz minus single pulse, 20 Hz minus 10 Hz, 40 Hz minus 10 Hz, and 40 Hz minus 20 Hz. For each pairwise contrast, the residual response was realigned to the first pulse that differed between the two protocols, thereby isolating the BF response attributed to that pulse. This procedure yielded six contrast-derived pulse-triggered responses. Together with the four original baseline-subtracted responses, these ten response estimates were averaged to generate one representative response trace per session. Session-level traces were then averaged within each animal.

This analysis assumes that pulse-triggered BF bursting has a common onset latency across stimulation protocols and pulse positions, although response amplitude may vary across conditions. Because the pooled trace combines responses derived from multiple stimulation protocols and pulse positions, the analysis was used specifically to estimate burst onset rather than to interpret the response waveform after onset. Burst onset latency was defined from the population response across animals as the first post-stimulation time bin in which the response became significantly greater than zero, assessed using a one-sample t test with Bonferroni correction for multiple comparisons across post-stimulation time bins. The first significant positive bin was taken as the earliest detectable BF burst onset following an Npr3 stimulation pulse.

#### Behavioral performance quantification

Behavioral performance in the auditory detection task was quantified by lick-response probability and reaction time across sound intensities. Lick-response probability, termed hit rate on sound trials, was defined as the proportion of trials in which mice produced three consecutive licks within the response window. Reaction time was defined as the latency from sound onset to the first lick.

#### Reward-prediction threshold model

To quantify the relationship between BF bursting activity and behavioral performance, we constructed a probabilistic threshold model that predicts behavioral response probability from trial-by-trial cue-period population BF burst amplitude. For each trial, BF burst amplitude, ΔFR, was calculated by averaging firing rates across simultaneously recorded BF bursting neurons within each session and subtracting the corresponding baseline firing rate. On each trial, the animal was assumed to compare ΔFR against an internal response threshold, T, drawn from a normal distribution:

T ∼ N(μ, σ²),

where μ represents the mean BF activity threshold for generating a behavioral response, and σ is the standard deviation of the threshold distribution, capturing trial-to-trial threshold variability. The behavioral response rule was defined as:

if ΔFR > T, the animal responds with probability 1 - λ;
if ΔFR ≤ T, the animal responds with probability κ.

Here, λ represents the upper lapse rate, and κ represents the baseline response rate, corresponding to the false-alarm rate in the absence of a stimulus. Marginalizing over the latent threshold distribution gives:

P(Hit) = κ + (1 - κ - λ) Φ((ΔFR - μ) / σ),

where Φ is the cumulative distribution function of the standard normal distribution.

Model parameters were estimated using non-stimulated sound-only trials alone. For each session, parameters were fitted by maximizing the likelihood of the observed behavioral outcomes (hit or miss) given the trial-by-trial BF burst amplitudes. The fitted model was then applied to Npr3-stimulated trials from the same session without refitting. This provided a test of whether the Npr3-induced increase in BF burst amplitude was sufficient to account for the stimulation-induced change in behavioral response probability. To quantify the predicted behavioral effect of Npr3 stimulation, we calculated the predicted gain as the difference in response probability between stimulated and sound-only trials. Predicted gains were compared with experimentally observed gains after averaging across sessions within each animal for each sound condition.

To generate model-predicted cue-period BF response patterns for hit and miss trials, we used the fitted probabilistic threshold model to simulate behavioral outcomes from the observed trial-by-trial BF burst amplitudes. For each trial, hit or miss outcomes were sampled stochastically according to the response probability determined by the fitted model parameters and the observed ΔFR. This simulation was repeated 1,000 times. For each simulation, PSTHs and burst amplitudes were averaged separately for predicted hit and miss trials, and the resulting averages were then averaged across simulations. This procedure tested whether the threshold model could reproduce the observed separation of cue-period BF responses between hit and miss trials, the increase in BF burst amplitude with sound intensity, and the upward shift induced by Npr3 stimulation (Figure S8).

#### Statistical analysis

Data are reported as mean ± s.e.m. unless otherwise indicated. For comparisons across more than two related conditions, Friedman tests were used, followed where appropriate by Bonferroni-corrected pairwise Wilcoxon signed-rank tests. Paired comparisons between matched conditions were assessed using Wilcoxon signed-rank tests. Statistical significance was defined as p < 0.05. Sample sizes and the corresponding units of analysis, including animals, sessions, neurons, or session-condition observations, are indicated in the figure legends and Table S1.

---

**Figure S1.**
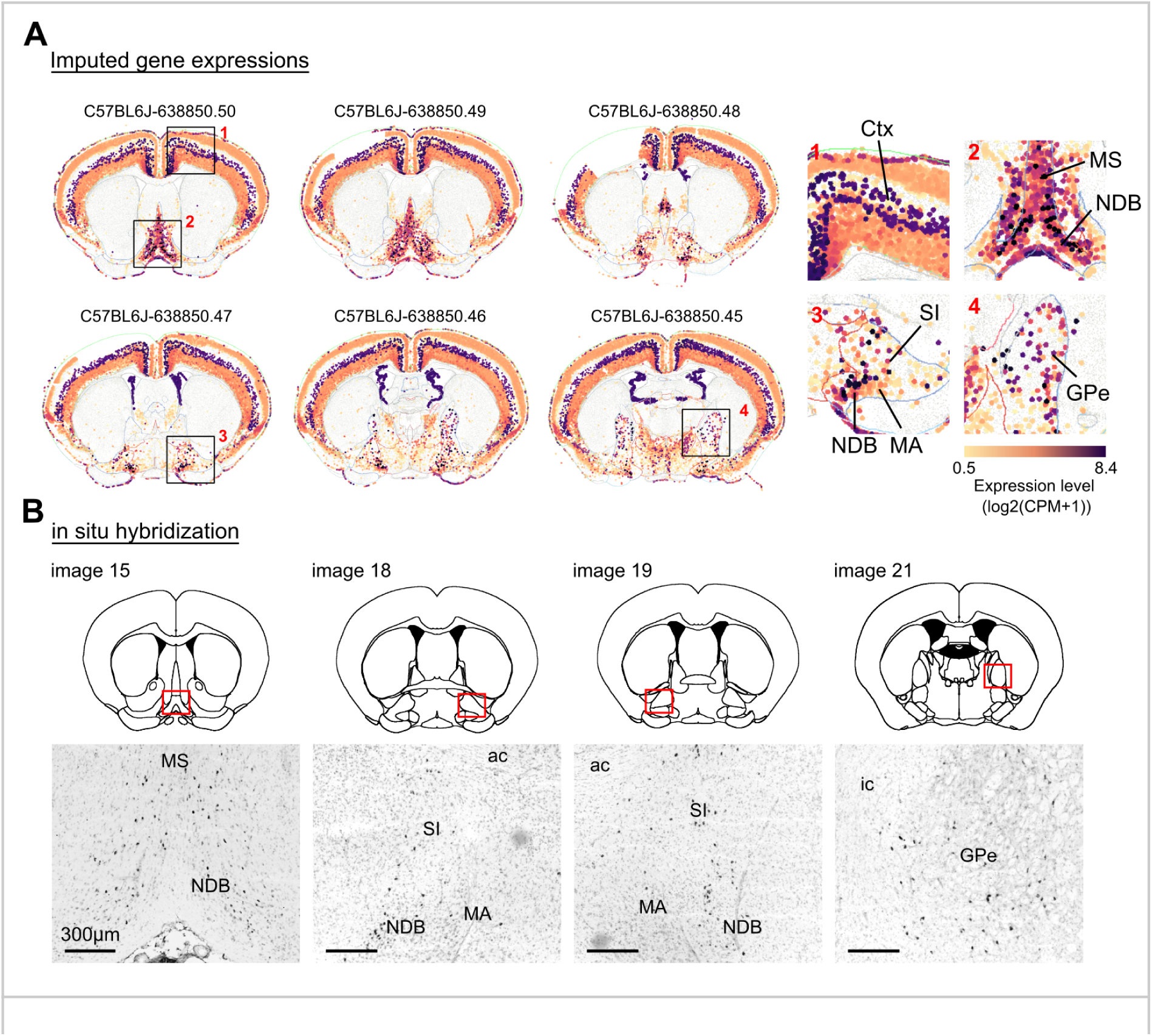
Npr3 expression follows the anatomical organization of the broader basal forebrain macrosystem. **A**, Spatial distribution of Npr3 expression visualized using the Allen Brain Cell Atlas, <u>MERFISH-C57BL6J-638850 with Imputed Genes dataset</u>^13^. While Npr3 is expressed abundantly in cortical neurons, its expression in subcortical regions is highly specific and concentrated in regions belonging to the broader basal forebrain macrosystem, including the medial septum (MS), nucleus of the diagonal band (NDB), substantia innominata (SI), magnocellular preoptic area (MA), and globus pallidus externus (GPe). Npr3 expression is sparse in surrounding subcortical regions, supporting its use as a candidate marker for a defined basal forebrain-associated population. **B**, Npr3 in situ hybridization from the Allen Mouse Brain Atlas (<u>Experiment ID</u> <u>73931412</u>)^41^, showing a similar regional expression pattern across basal forebrain-related territories.

**Figure S2.**
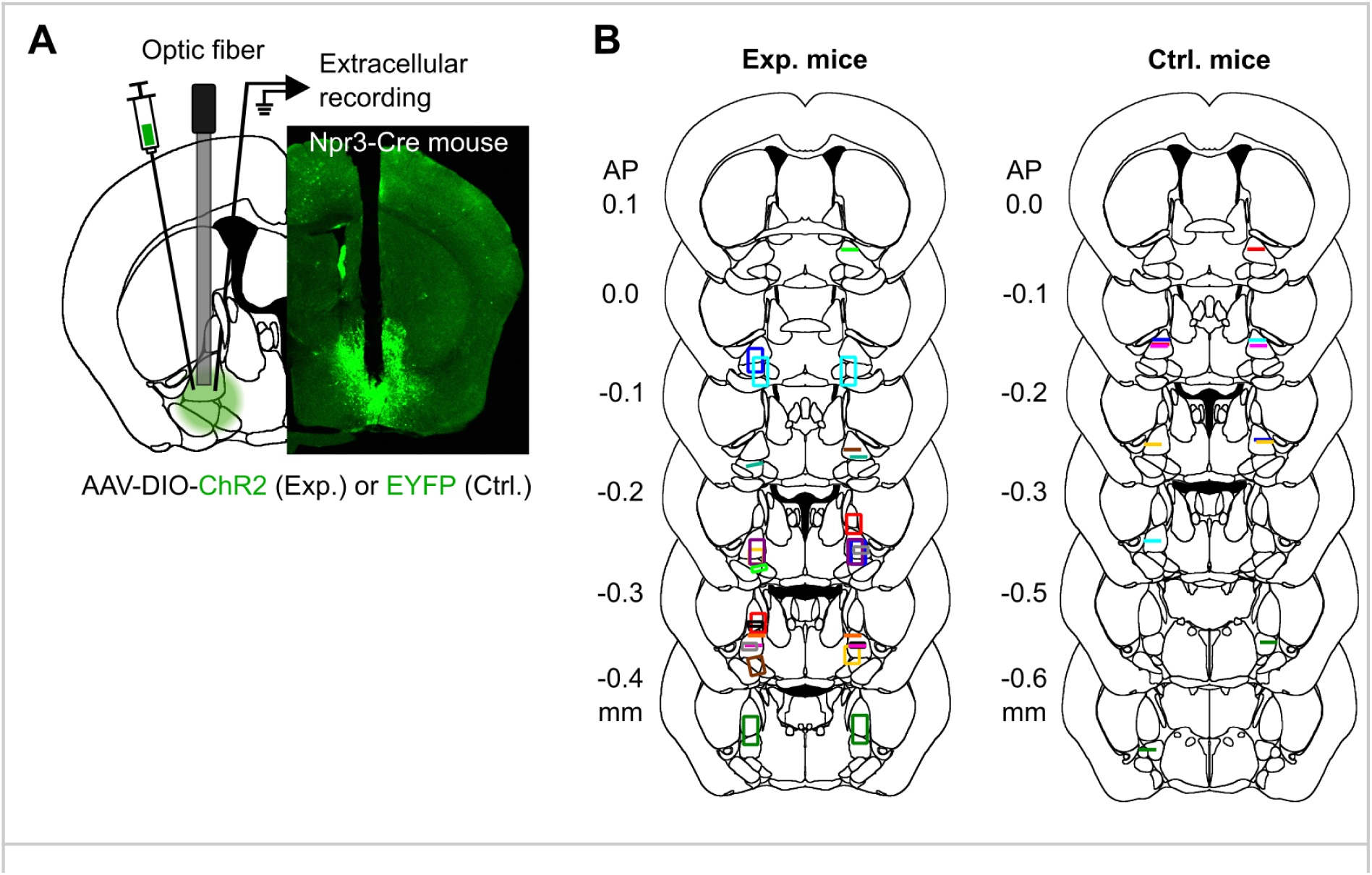
Histological verification of optic fiber placement and BF recording sites. **A**, Schematic of the experimental configuration for optogenetic stimulation and extracellular recording of BF Npr3 neurons. Npr3-Cre mice received bilateral BF injections of AAV-DIO-ChR2; control mice received AAV-DIO-EYFP. In most experimental mice, bilateral microdrives were implanted with an optic fiber and tungsten electrodes bundled within each cannula, allowing electrodes to be advanced across recording sessions. Right, representative ChR2-EYFP expression in the BF together with the cannula tract. **B**, Histological reconstruction of optic fiber placements and BF recording sites across coronal levels in experimental (N=13) and control mice (N=6). Colored symbols denote individual animals. Line segments indicate optic fiber locations in mice implanted with optic fibers alone, including control mice and experimental mice without electrophysiological recordings. Rectangles indicate the recording regions sampled by microdrive implants; their dorsoventral extent represents the range of electrode depths sampled across recording sessions. Anteroposterior coordinates are given relative to bregma. Coronal atlas outlines were adapted from the Allen Mouse Brain Common Coordinate Framework (CCFv3)^42^.

**Figure S3.**
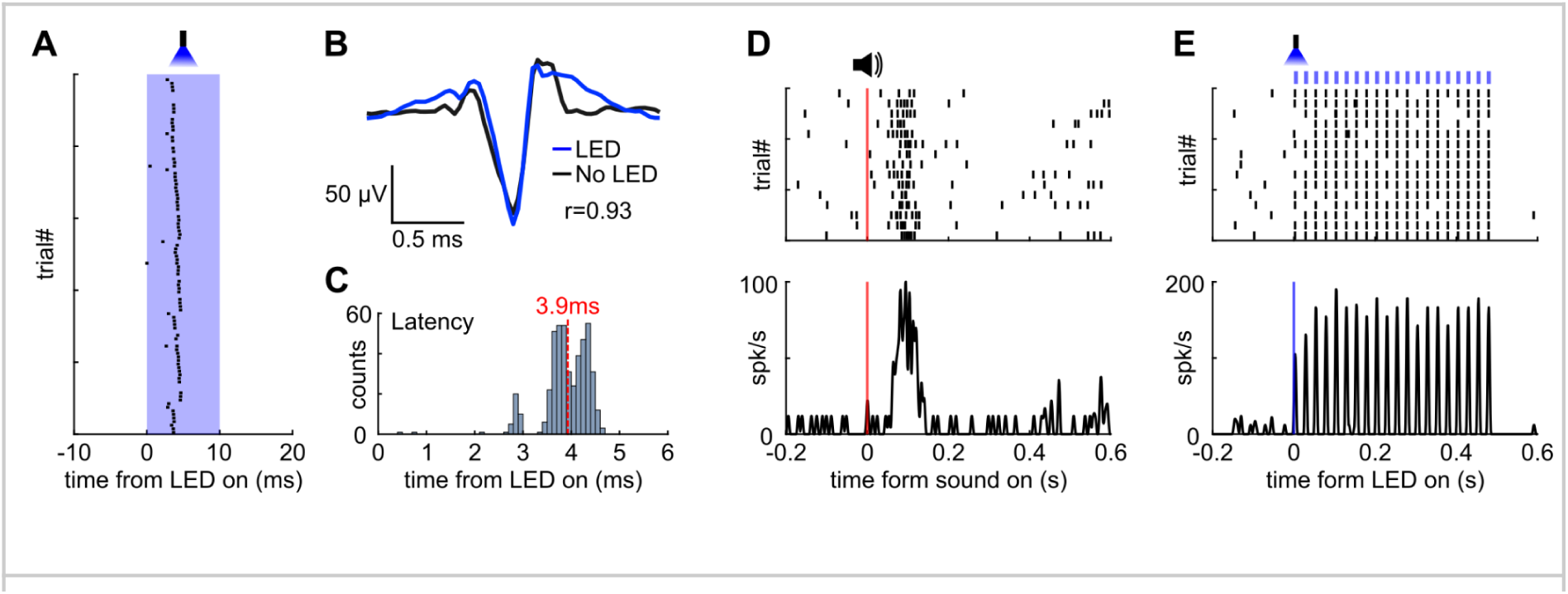
Example phototagged Npr3 neuron recorded in the basal forebrain. **A**, Spike raster of an example phototagged Npr3 neuron aligned to optogenetic stimulation. **B**, Average spike waveform. Blue: average light evoked spikes; black: average spontaneous spikes (Pearson correlation, r = 0.93). **C**, Spike latency distribution after optogenetic stimulation. Red dashed line, mean. **D**, Response of the phototagged Npr3 neuron to the reward-predicting sound during the auditory task (Figure 1E). The response pattern resembles a typical BF bursting neuron. **E**, Phototagged Npr3 neuron aligned to 40Hz optogenetic stimulation. Note that phototagged Npr3 neurons were rare in our recordings, precluding population-level analysis; this neuron is shown as an illustrative example.

**Figure S4.**
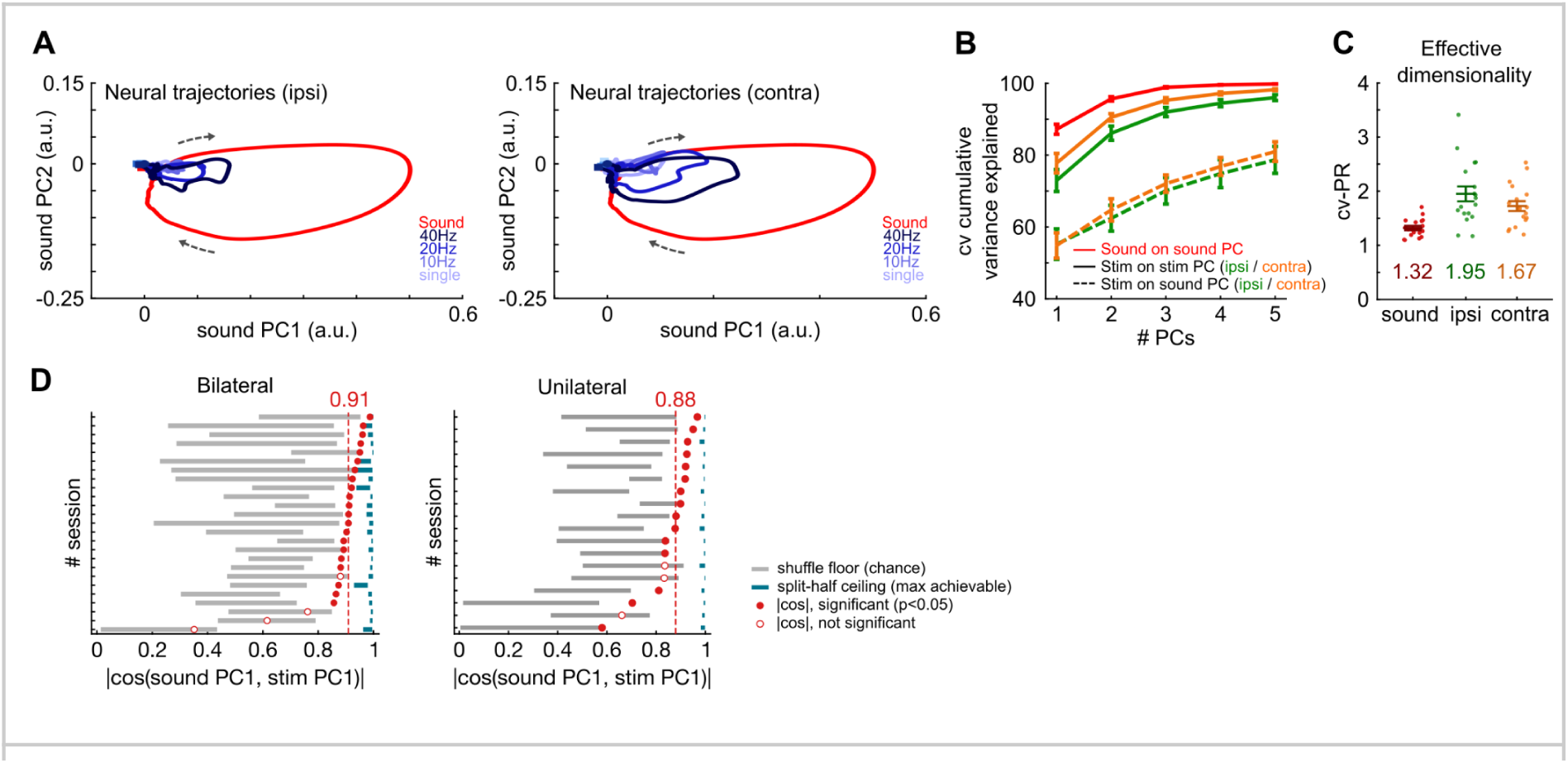
Unilateral Npr3 stimulation recruits the low-dimensional population dynamics of natural BF bursting. **A**, Population neural trajectories, using the same analysis as in Figure 1Q and Figure 2F, shown separately for ipsilateral (left) and contralateral (right) Npr3 stimulation. Mean sound responses (red) and stimulation responses (blue) were projected onto the first two PCs defined by the sound response. Arrows indicate progression over time from baseline through the burst and back to baseline. **B**, Cross-validated cumulative variance explained by the first k PCs, using the same analysis as in Figure 1R and Figure 2G. Sound responses projected onto sound PCs are shown in red. Ipsilateral (green) and contralateral (orange) stimulation are shown projected onto their own PCs (solid lines; ceiling) and onto the sound PCs (dashed lines; alignment). Mean ± s.e.m. across 18 sessions. **C**, Effective dimensionality, quantified by cross-validated participation ratio (CV PR), as in Figure 1R and Figure 2G, for sound (red), ipsilateral stimulation (green), and contralateral stimulation (orange). Dots indicate individual sessions; symbols and error bars indicate mean ± s.e.m., with group means shown below. **D**, PC1 alignment, quantified as |cos θ| between sound and stimulation PC1 loading vectors (1, identical; 0, orthogonal), using the same analysis as in Figure 1R but shown separately for each session to compare the observed alignment with the expected floor and ceiling. Bilateral stimulation is shown at left for comparison with unilateral stimulation at right. Each row represents one session, sorted by |cos θ|. Gray bars indicate the neuron-identity shuffle floor, and green bars indicate the sound split-half ceiling, representing the best achievable alignment at the matched trial count. Filled dots indicate significant alignment relative to the shuffle distribution (p < 0.05); open dots indicate non-significant sessions. Red dashed lines indicate the median |cos θ| across sessions.

**Figure S5.**
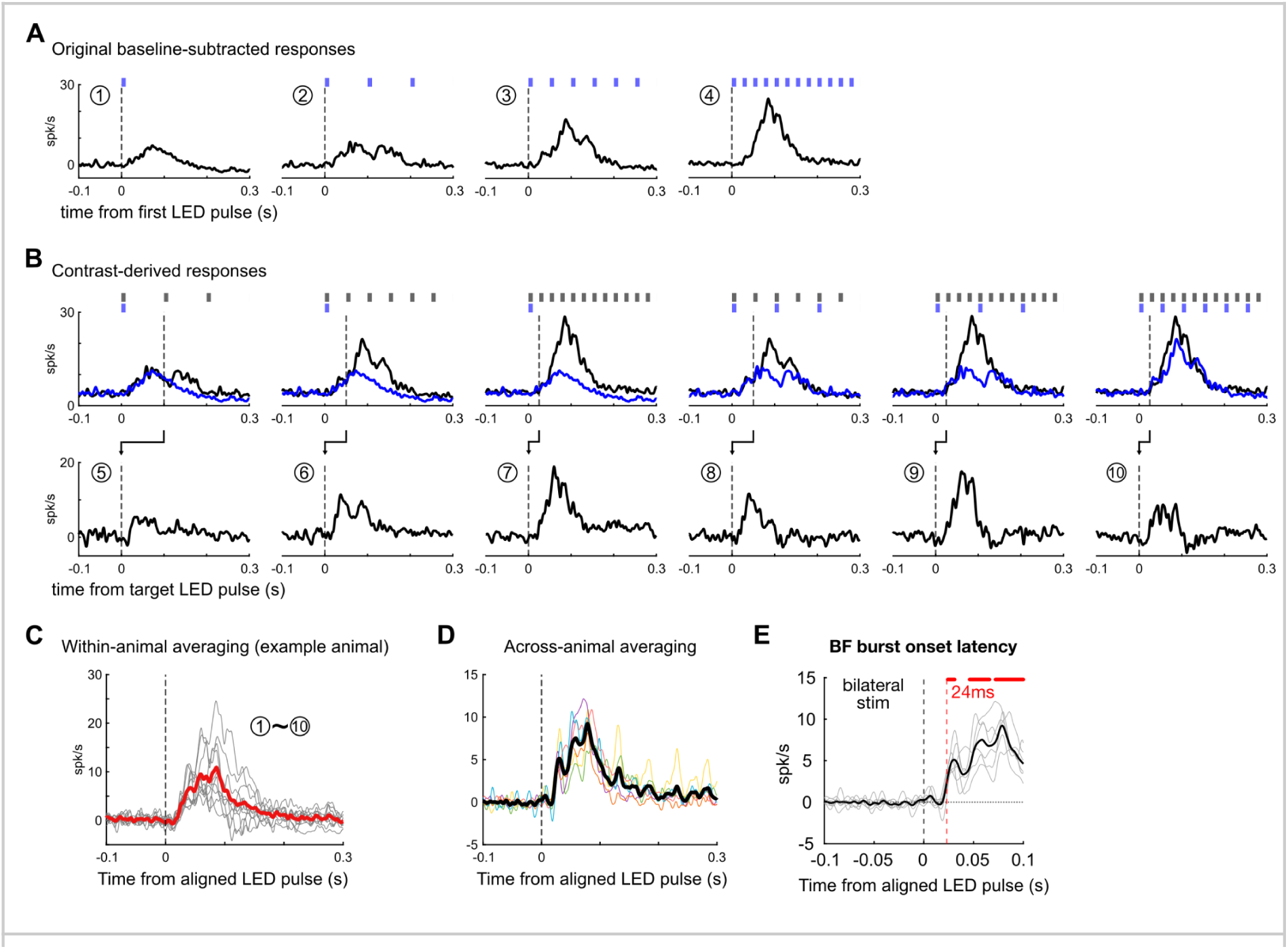
Estimation of BF burst onset latency following single-pulse Npr3 stimulation. Panels A-D illustrate the latency-estimation procedure using bilateral Npr3 stimulation (n = 6 mice), with the resulting onset estimate shown in E. The same procedure was applied separately to ipsilateral and contralateral stimulation to obtain the estimates shown in Figure 2H. To obtain a robust estimate of BF burst onset latency following an Npr3 stimulation pulse, we combined responses to the first pulse of the original stimulation protocols with contrast-derived responses that isolated responses to later pulses within stimulation trains. **A**, Original baseline-subtracted population responses to single-pulse, 10-Hz, 20-Hz and 40-Hz Npr3 stimulation, aligned to the first LED pulse. **B**, Six pairwise contrasts used to derive additional pulse-triggered responses. Upper plots show the two responses entering each subtraction (black minus blue), with the corresponding stimulation patterns above. Lower plots show the resulting difference traces, realigned to the target LED pulse, defined as the first pulse at which the two stimulation protocols differ. **C**, Response estimates generated from the original and contrast-derived responses were aligned to stimulation onset and averaged to obtain a single latency-estimation trace for each animal. The example shows the responses contributing to the average for one animal. **D**, Animal-level estimates were combined to generate the population response used to determine BF burst onset latency. Because this pooled response combines estimates derived from multiple stimulation protocols and pulse positions, it is intended primarily to estimate the earliest detectable burst onset; the response waveform following onset is not interpreted as the response to any individual stimulation protocol. **E**, Estimated BF burst onset latency following bilateral Npr3 stimulation. Gray lines show individual-animal traces (n = 6 mice), and the black line shows the population mean. The red dashed line marks the estimated onset at 24 ms, defined as the first post-stimulation time bin with a population response significantly greater than zero.

**Figure S6.**
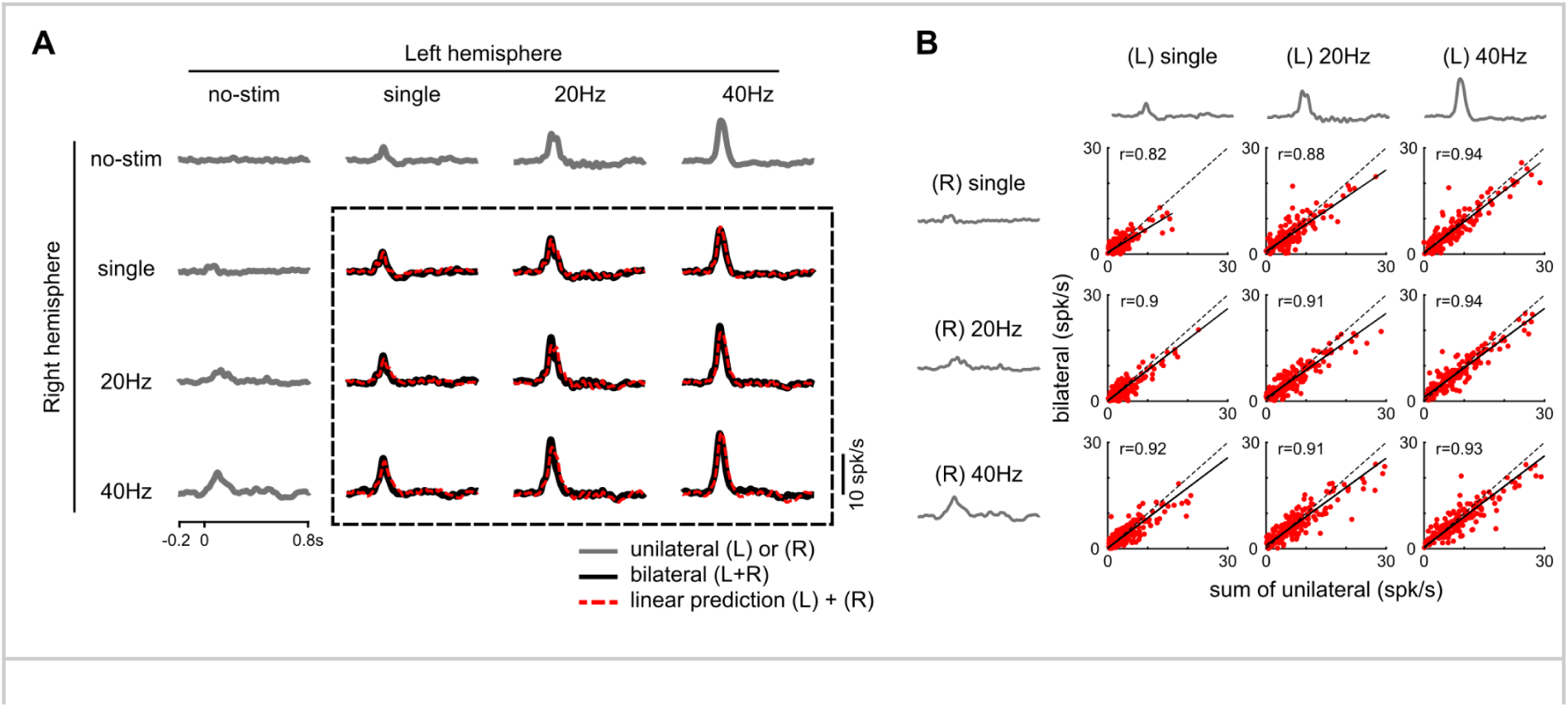
Bilateral BF bursting responses are approximated by summed unilateral responses across sessions and neurons. **A**, Extension of the analysis shown in Figure 3C to all recorded sessions. Left and right BF hemispheres were stimulated independently at four levels: no stimulation, single pulse, 20 Hz, and 40 Hz, yielding 16 stimulation combinations. Population-average PSTHs are plotted using the same conventions as in Figure 3C. Gray traces outside the dashed rectangle show measured responses to unilateral or no-stimulation conditions. Black traces inside the dashed rectangle show measured responses to the nine bilateral stimulation combinations, and cyan traces show the corresponding summed-unilateral predictions. **B**, Extension of the measured-versus-predicted comparison shown in Figure 3D to individual BF bursting neurons. For each of the nine bilateral stimulation combinations outlined in A, measured bilateral responses are plotted against the corresponding summed-unilateral predictions. Each dot represents one neuron (n=202). Dashed lines indicate unity, and solid lines indicate linear fits. Across stimulation combinations, bilateral responses closely followed the summed-unilateral predictions, supporting approximate linear summation at the single-neuron level.

**Figure S7.**
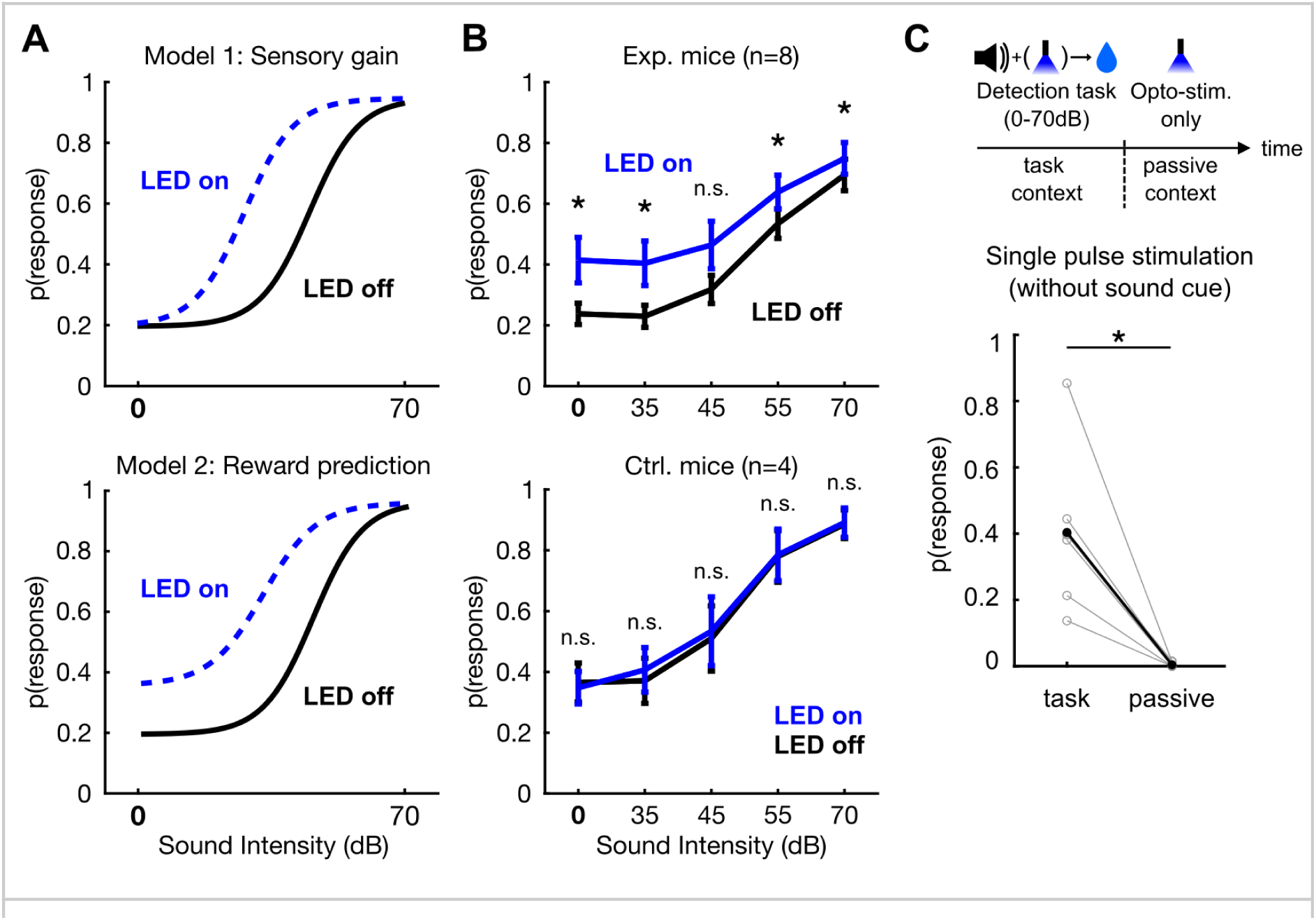
Behavioral enhancement by Npr3 stimulation is not explained by sensory gain or direct motor activation. **A**, Two alternative models for how Npr3 activation could improve detection performance. Under a sensory-gain model, Npr3 activation would shift the psychometric curve leftward by increasing effective sound intensity. Under a reward-prediction model, Npr3 activation would increase response probability even when no sound stimulus is presented. **B**, Test of model predictions using sessions in which the lowest sound level was replaced by a no-sound condition (0 dB). In experimental mice, Npr3 stimulation increased response probability even in the 0 dB condition. This effect was absent in EYFP control mice (Bonferroni-corrected Wilcoxon signed-rank tests; n = 8 experimental mice and 4 control mice; *p < 0.05). **C**, Test of whether Npr3 stimulation directly triggered licking. During the behavioral task, single-pulse Npr3 stimulation in 0 dB trials induced licking despite the absence of a sound cue. In contrast, the same stimulation delivered passively outside the task evoked little licking. Stimulation-associated response probability was substantially lower in the passive context than during task performance, arguing against a direct motor effect of Npr3 stimulation (Wilcoxon signed-rank test; n = 7 animals; *p < 0.05).

**Figure S8.**
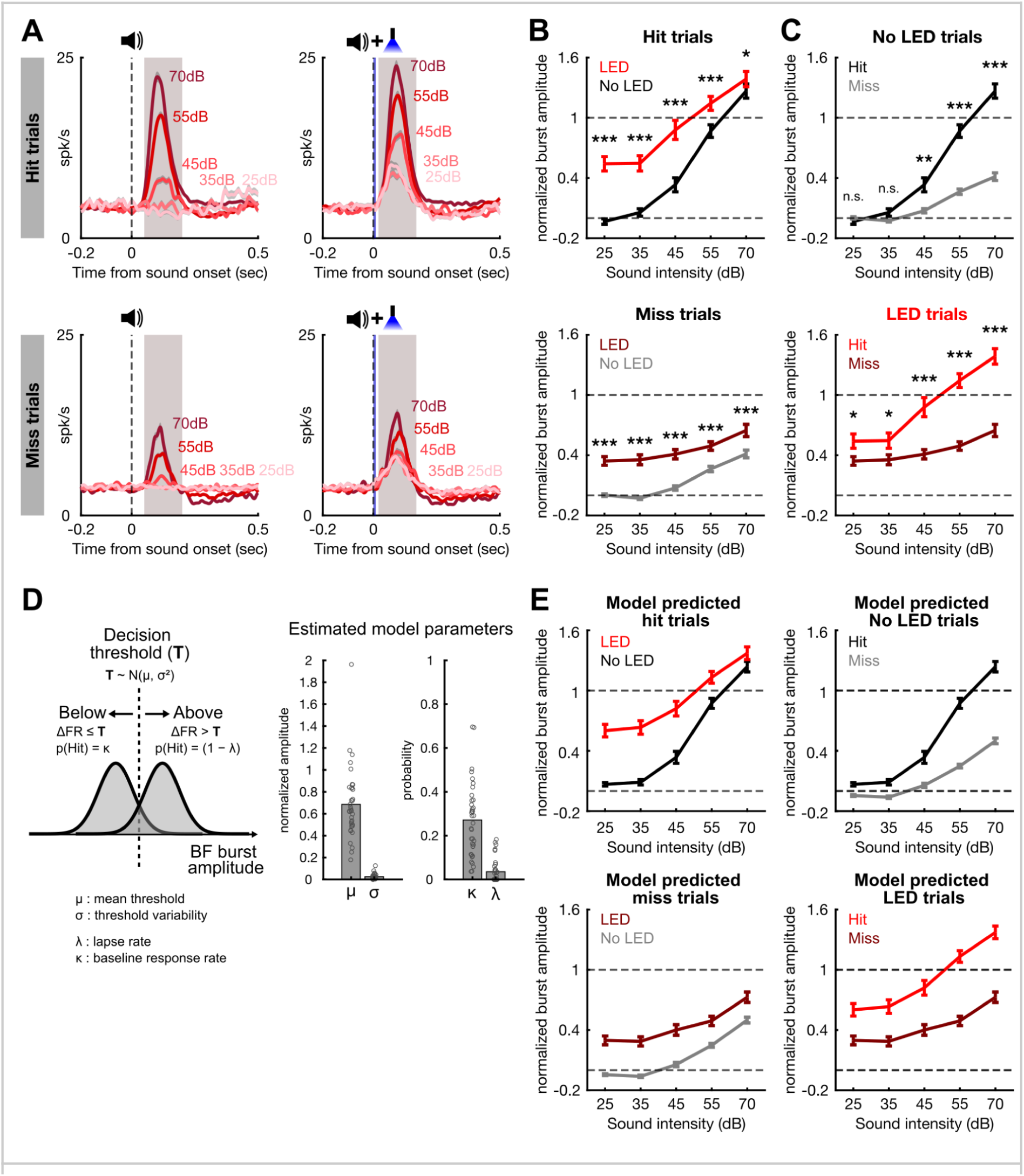
Threshold-model analysis of cue-period BF bursting responses in hit and miss trials. **A**, Cue-period BF bursting responses in hit and miss trials, separated by no-LED and LED trials. PSTHs show population responses of BF bursting neurons aligned to sound onset. Responses are shown separately for hit trials and miss trials across sound intensities. Shaded areas indicate the analysis window used to quantify cue-period BF burst amplitude. **B**, Summary of normalized cue-period BF burst amplitudes in hit trials and miss trials, comparing LED and no-LED conditions across sound intensities. Npr3 stimulation shifted BF burst amplitudes upward in both hit and miss trials. Corresponding LED and no-LED conditions were compared at each sound intensity using Bonferroni-corrected Wilcoxon signed-rank tests (n = 36 sessions; n.s., not significant; *p < 0.05, **p < 0.01, ***p < 0.001). **C**, Observed cue-period BF burst amplitudes in hit and miss trials, shown separately for no-LED and LED trials. In both conditions, hit trials showed larger BF burst amplitudes than miss trials, and BF burst amplitude increased with sound intensity. Hit and miss trials were compared at each sound intensity using Bonferroni-corrected Wilcoxon signed-rank tests (n = 36 sessions; n.s., not significant; *p < 0.05, **p < 0.01, ***p < 0.001). **D**, Schematic and fitted parameters of the probabilistic threshold model. The model assumes that behavioral responses are generated by comparing cue-period BF burst amplitude, ΔFR, with an internal decision threshold, T, drawn from a normal distribution, T ∼ N(μ, σ²). When ΔFR exceeds T, the animal responds with probability 1 - λ; when ΔFR does not exceed T, the animal responds with probability κ. μ represents the mean decision threshold, σ represents threshold variability, λ represents the lapse rate, and κ represents the baseline response probability. Bar plots show fitted model parameters across sessions. μ and σ are expressed in normalized BF burst-amplitude units, using the same normalization as in Figure 3G. Each dot represents the estimate from one session (n = 36 sessions). **E**, Model-predicted cue-period BF burst amplitudes in hit and miss trials. Behavioral outcomes were simulated from the fitted probabilistic threshold model using the observed trial-by-trial BF burst amplitudes. For each trial, hit or miss assignment was sampled stochastically from the response probability determined by the fitted model parameters and the observed ΔFR. This simulation was repeated 1,000 times, and PSTHs and burst amplitudes were averaged separately for predicted hit and miss trials across simulations. The model reproduced the higher BF burst amplitudes in hit trials, the lower amplitudes in miss trials, the increase in burst amplitude with sound intensity, and the upward shift induced by Npr3 stimulation.

**Table S1.** Summary of animals and sessions contributing to each experiment and analysis. Numbers of animals and recording or behavioral sessions included in each experimental group, together with the corresponding main and supplementary figures. Animal numbers are separated by sex. For the experimental group in the 25–70 dB detection task, recording (rec.) and behavioral (beh.) sessions are reported separately.

| **Experiment** | **Animals / Sessions** | **Figure** |
| --- | --- | --- |
| Bilateral stimulation | 6 male / 25 sessions | Fig. 1, S4, S5 |
| Unilateral stimulation | 5 male / 19 sessions | Fig. 2, S3, S4 |
| Bilateral input summation | 2 male, 1 female / 7 sessions | Fig. 3, S6 |
| Detection task -<br>exp. group (25-70dB) | 9 male / 36 sessions (rec.)<br>9 male, 1 female / 200 sessions (beh.) | Fig. 3, 4, S8 |
| Detection task -<br>ctrl. group (25-70dB) | 4 male, 2 female / 130 sessions | Fig. 4 |
| Detection task -<br>exp. group (0-70dB) | 6 male, 2 female / 100 sessions | Fig. S7 |
| Detection task -<br>ctrl. group (0-70dB) | 2 male, 2 female / 66 sessions | Fig. S7 |

## Acknowledgements

We thank T.-W. Chen for critical discussions of the manuscript; S.-T. Wu for technical support.

## Funding

This research was funded by the Ministry of Science and Technology, Taiwan (MOST 108-2638-B-010-002-MY2 to S.-C.L.); National Science and Technology Council, Taiwan (NSTC 110-2628-B-A49A-505, 111-2628-B-A49-015, 112-2628-B-A49-013, 113-2628-B-A49-001, 114-2320-B-A49-029-MY3 to S.-C.L.); and from the Brain Research Center, National Yang Ming Chiao Tung University through The Featured Areas Research Center Program within the framework of the Higher Education Sprout Project by the Ministry of Education (MOE) in Taiwan.

## Contributions

H.-C.L. and S.-C.L. conceived and designed the study. H.-C.L. performed the experiments. H.-C.L. and S.-C.L. analyzed the data and wrote the manuscript. S.-C.L. supervised the research.

## Competing interests

Authors declare that they have no competing interests.

## Notes

### Competing Interest Statement

The authors have declared no competing interest.

